# Internal Normal Mode Analysis applied to RNA flexibility and conformational changes

**DOI:** 10.1101/2022.11.30.518608

**Authors:** Afra Sabei, Talissa Gabriele Caldas Baia, Raphaël Saffar, Juliette Martin, Elisa Frezza

**Affiliations:** Université Paris Cité, CiTCoM, CNRS, F-75006 Paris, France; Univ Lyon, Université Claude Bernard Lyon 1, CNRS, UMR 5086 MMSB, Lyon, France

## Abstract

We investigated the capability of internal normal modes to reproduce RNA dynamics and predict observed RNA conformational changes, and, notably, those induced by the formation of RNA-protein and RNA-ligand complexes. Here, we extended our iNMA approach developed for proteins to study RNA molecules using a simplified representation of RNA structure and its potential energy. Three datasets were also created to investigate different aspects. Despite all the approximations, our study shows that iNMA is a suitable method to take into account RNA flexibility and describe its conformational changes opening the route to its applicability in any integrative approach where these properties are crucial.

## Introduction

Ribonucleic acid (RNA) molecules are involved in nearly any physiological phenomenon in the cell.^1–3^ Moreover, many viruses (e.g., HIV, Hepatitis C, coronavirus) manipulate cellular machineries to ensure their replication, for instance to translate their mRNA.^4^ RNA functions crucially rely on both the specific three-dimensional (3D) folding of the molecule, which in turn depends on the sequence and on how nucleobases pair through hydrogen bonds,^5^ and its conformation. This relationship is even more crucial for protein-RNA complexes. The dysfunction of such complexes is implicated in many humans and animal pathologies.^6^ For example, the formation of ribonucleoprotein particles (RNPs) including mRNAs is key for the post-transcriptional regulation of gene expression. ^7^ As for proteins, RNA molecules can undergo conformational changes based on different stimuli (for example pH) or following the binding to a ligand or another biomolecule that often goes beyond local rearrangements.^8–10^ A well-known example is given by the SAM-riboswitch which can change its 2D folding to bind the *S* -adenosyl-*L*-methionine.^11^ Hence, RNA flexibility is crucial for its function and underlies not only RNA folding, but also the majority of RNA interactions with other molecular species.^12^ However, its flexibility is very complex allowing to adopt distinct conformations and it is one of the major reasons why obtaining high-resolution 3D structures via X-ray crystallography, NMR or cryo-electron microscopy (cryoEM) is still a challenging task as shown by the relatively small number of bound or unbound structures deposited in the Nucleic Acid Data Bank (NDB).^13,14^ In this context, predicting both the structure and structural transitions and conformational changes linked to the interactions is very important, particularly in view of the increasing use of relatively low-resolution data obtained by small-angle X-ray scattering (SAXS), cryoEM or biochemical probing (for example Selective 2’-Hydroxyl Acylation analyzed by Primer Extension, SHAPE)^15–17^ and to develop new integrative approaches.^18–24^

Predicting local or global changes in RNA conformations is still very difficult. In some cases, this hinders the prediction of the structure of the complex or the change of RNA folding.^12^ A common method to study RNA flexibility, RNA dynamics and conformational rearrangements for isolated RNA molecules or RNA-ligand/RNA-protein complexes are molecular dynamics simulations using either an all-atom or a coarse-grain representation. However, the millisecond simulations currently reachable^25^ are computationally expensive, several replica are needed and enhanced methods are still limited for these tasks and the size of RNA molecules is small.^12,22,26–29^ Moreover, these methods are still of limited use for RNA-protein/RNA-ligand docking without reliable starting conformations. Therefore, simplified and faster methods are highly sought. Among them, normal mode analysis (NMA)^30^ is a well-known computational approach to represent the intrinsic flexibility of biomolecules and their conformational changes and it is typically combined with coarse-grain molecular representations and simplified energy models, like elastic network models (ENM) and in particular Anisotropic Network models (ANM).^26,31–37^

Under a harmonic approximation, NMA gives insights into the equilibrium vibrational modes accessible to a system. After many decades of applications to classical physical problems, molecular applications to biomolecules, in particular proteins, confirmed the importance of low-frequency motions in biological processes.^30,38,39^ Although NMA has largely been applied to study proteins and protein deformations induced by ligand binding, allosteric processes and conformational changes occurring between isolated proteins and their complexes,^30,40–43^ the use of NMA for nucleic acids is still quite limited, in particular for single-stranded RNA. For example, NMA has been used in combination with experimental data such as SAXS^44,45^ and SHAPE^26^ to take into account RNA flexibility and its structural rearrangements. NMA has also been employed to study the dynamics of binding interactions of snRNA and U1A.^33^ Van Wynsberghe and Cui investigated the hammerhead ribozyme and a guanine riboswitch and have shown that NMA coupled with ENM models can reproduce key aspects of nucleotide dynamics but that it may not be as precise for loosely packed structures as it is for densely packed ones, such as globular proteins.^46^ In this perspective, Zimmermann and Jernigan have investigated the capability of elastic network models and coarse-grain models coupled with NMA to capture the apparent motions within ensembles of 16 RNA structures.^32^ Their study shows that ENM models and NMA are suitable methods to study well-packed RNA-only structures, thus justifying their use in the analysis of the dynamics of protein-RNA complexes such as those involving ribonucleic proteins.

As for proteins, almost all the studies reported in the literature for ssRNA are referring to NMA computed using a Cartesian Coordinate Space (CCS) for its simplicity (hereafter termed cNMA). This approach is often combined with a coarse-grain model based on a simplified RNA backbone (i.e., the Cartesian positions of the P atoms) where the pseudoatoms are connected by linear springs within a given cut-off distance or more complex dependencies.^32,33^ More structural information on both the backbone and the nucleobase atoms can also be included.^26^ However, cNMA suffers from the fact that the harmonic approximation is only valid for relatively small movements, implying that conformational changes defined by the normal modes will quickly deform the valence structure (i.e., bond lengths and bond angles) of RNA. These intrinsic limitations can explain why cNMA fails to correctly predict the dynamics of some RNA molecules, in particular for loosely packed structures, and the need to refine the structure if RNA undergoes large conformational changes.^46^

One clear route to improve NMA involves the choice of another coordinate space. Gō and co-workers proposed an alternative approach based on the Internal Coordinate Space (ICS). They noted that ICS (torsion angles, bond angles, and bond lengths) is more advantageous as it extends the validity of the harmonic approximation of the conformational energy hypersurface.^47^ Thus, larger conformational changes can be modeled by taking into account the eigenvectors of low-frequency modes. This approach also allows to chose the “chemically relevant” variables, thus making it easier to split the degrees of freedom into two categories: “hard” (typically, bond lengths and valence angles) and “soft” (torsion angles). Based on the question, the hard variables can be excluded since they are unlikely to significantly contribute to medium/large collective movements, thus simplifying and speeding up the computations. For the reasons above, ICS NMA (hereafter named iNMA) is usually performed in torsional angle space, while keeping fixed valence angles and bond lengths, allowing thus to greatly reduce the total number of variables involved. As a consequence, an iNMA analysis can be carried out on any relevant subset of variables without modifying the molecular representation of the RNA molecule under investigation. Therefore, this approach allows to collect important pieces of information regarding those variables responsible for low-frequency collective movements. Moreover, iNMA, as cNMA, can equally describe spatial movements by a conversion of the modes into Cartesian space.^31,48^

Despite these advantages, applications of iNMA are relatively scarce in particular for ssRNA,^49^ probably due to the higher complexity behind it. This complexity comes from the fact that, in ICS, internal variables must be separated by overall rotations or translations of the molecular system under study. To do so, we need to know the topology of the system under investigation, in particular which pseudoatoms are moved by any given variable.^50–52^ Moreover, although iNMA represents a suitable tool for the characterization of the variables responsible for low-frequency collective movements, a direct description of the movements occurring in Cartesian space is not provided. To solve this problem, a second-order expansion can be used allowing to combine the advantages of iNMA with a CCS description of the overall conformational change, as shown for proteins in previous works.^31,48,53^

In the present work, we will use the iNMA approach (coupled with a simplified RNA representation and an elastic network energy model) to investigate the ability of internal normal modes analysis to capture RNA dynamics within 21 different ensembles of experimentally determined RNA structures and that obtained by all-atom molecular dynamics (MD) simulations for 16 RNA molecules. Finally, we will also study the unbound-to-bound transition for a set of RNA molecules for which at least two distinct conformational states are known and conformational changes are involved: an isolated, unbound state and a bound state involving interactions with another biomolecule or a ligand. In our framework, in order to extract the dynamics observed in the experimental ensembles and obtained by MD simulations, we will perform a Principal Component Analysis (PCA) in Cartesian space. Hence, to compare both RNA dynamics with the one predicted by iNMA and the experimental structures with those predicted by iNMA, we will do a conversion from the torsional space into Cartesian space without using any information from the target structure, as proposed earlier by Bray and co-workers,^54^ either using the second-order expansion for the first time applied to RNA molecules or the approach proposed by some of us in.^53^ The main aim of this work is to test the possibility of using low-frequency modes to describe RNA dynamics and unbound-to-bound RNA conformational transformations without introducing any deformation in the valence structure (bond lengths or angles) as would be the case with cNMA. We also aim to assess how the topology influences our results and if our approach is more appropriate to study loosely packed structures than cNMA. These aspects are important from the perspective of using iNMA in docking algorithms or in hybrid approaches combining NMA with experimental data from multiple sources to generate high-resolution models of RNA and RNA-biomolecules or RNA-ligand assemblies.

## Internal Normal Mode Analysis

### General Theory

Normal modes^55^ present an analytical solution of the classical equations of motion by imposing a harmonic approximation on the potential energy of the system (i.e., by assuming that the energy is a quadratic function of its N coordinates) around the potential energy minimum 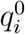

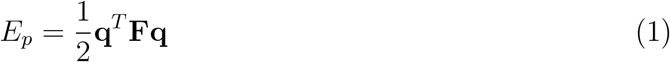

where **F** is the Hessian matrix, or potential energy matrix, defined by 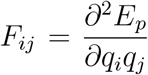 with *q*_*i*_ and *q*_*j*_ the coordinate *i* and *j*, and **q** is a set of internal coordinates. The kinetic energy of the molecule, *E*_*k*_, can then be expressed in terms of internal variables

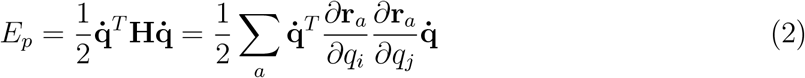

where **r**_*a*_ and m_*a*_ represent the position vectors and the atomic masses of each atom *a*, and 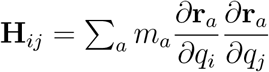 is the kinetic matrix. The internal coordinates **q** can be bond lengths, valence angles, torsion or rigid-body movements (rotations and translations). The equations of motion in terms of any set of coordinates for *n* variables are given by Lagrange’s equations, whose solution takes the following form

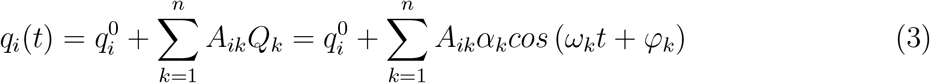

where *α*_*k*_ and *φ*_*k*_ depend on the initial conditions and are the thermal amplitude and the phase of the *k*-th mode, respectively. The unknowns, *A*_*ik*_ and *ω*_*k*_, are obtained by solving the generalized eigenvector problem:

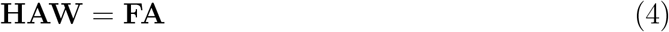

where **A** is the matrix of the eigenvectors and **W** is a diagonal positive matrix whose elements are

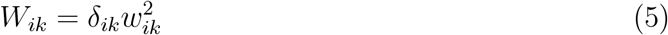

with *δ*_*ik*_ the Kronecker delta.

The amplitude *α*_*k*_ of a given normal mode *k* is also called thermal amplitude and it depends on the temperature. Using the equipartition theorem, each normal mode has a time-averaged potential energy equal to 1/2*k*_*B*_*T*, with *T* the absolute temperature and *k*_*B*_ the Boltzmann constant. Since the time-averaged potential energy of each mode can be expressed as

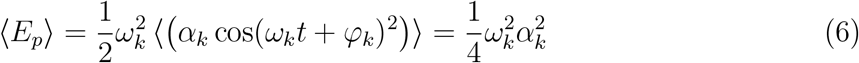

we can obtain the thermal amplitude

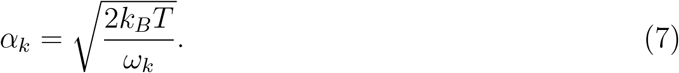

where *ω*_*k*_ is the frequency of the kth mode.

### Internal Coordinate Normal Analysis

In the presence of two RNA strands or another biomolecule, two types of variables must be taken into account: interbody and intrabody coordinates. The number of independent interbody variables is 6*N* -6, with *N* being the number of independent strands or molecules. For each body, we chose the P atom for RNA molecules and C*α* atom for proteins^31^ closest to the center of mass as the pivot for the rotation and translation. A complete description of the variables can be found in our previous work. ^31^

#### Hessian matrix

To compute the Hessian matrix, for the first derivative we used an analytical calculation as in our previous work.^31^ The derivative of the energy with respect to the independent internal variables *q*_*i*_ can be written as:

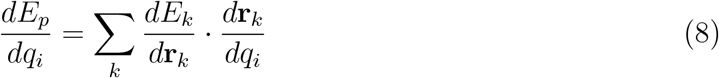

where the sum runs over the subset of atoms *k* moving when a rotation is made around the unit bond vector **b**_*i*_ = (**r**_*i*+1_ − **r**_*i*_)/*s*_*i*_. For the torsional angle *τ*_*i*_, *s*_*i*_ = |**r**_*i*+1_ − **r**_*i*_| and 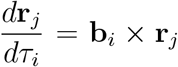 with **R**_*ji*_= **r**_*j*_ − **r**_*i*_. In the case of the valence angles, *d***r**_*j*_ /*dθ*_*I*_ = (**R**_*ji*_ × **c**_*i*_), where **c**_1_ = (**b**_*i-*1_ − **b**_*i*_)/ sin *θ*_*i*_ is the perpendicular vector to the plane of the valence angle; in the case of the distances, *d***r**_*j*_/*dt*_*i*_ = **b**_*i*_. The second derivatives are computed numerically for computational efficiency as presented by some of us in.^31^

#### Kinetic matrix

Regarding the kinetic matrix, **H**, the derivatives of the atomic position vector **r**_*a*_ with respect to the independent variables *q*_*i*_ must be computed carefully. In fact, as the internal kinetic energy cannot include external motions (i.e., the overall translation and rotation of the system), the change *δ***r**_*a*_ caused by *δq*_*i*_ must not move the center of mass or modify the inertia tensor. To this end, Noguti and Gō proposed an elegant analytical approach to compute **H** in dihedral angular space (DAS).^50,56^ In other words, for a given dihedral angle, taking all the other variables as fixed, the system can be treated as two rigid bodies connected by a chemical bond around which a rotation *τ*_*p*_ can occur. For each atom b in the first body and each atom c in the second, *δ***r**_*b*_ and *δ***r**_*c*_ are expressed using Eckart conditions to separate internal and external motion.^57^ All the details can be found in a previous publication by some of the present authors.^31^

### Conversion from Internal Coordinates to Cartesian Ones

For small-amplitude conformational dynamics of RNA, the Taylor expansion of the Cartesian coordinates^31,48,53^ around a given conformation (usually an energy minimum) corresponding to a set of internal coordinates **q** is given by

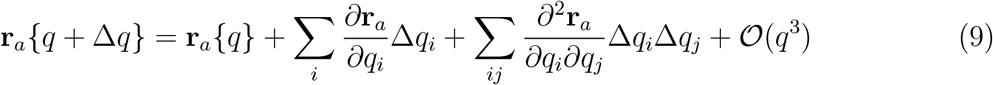

where Δ*q*_*i*_ is the displacement of the internal coordinate *i* in the preceding instantaneous conformation, defined as

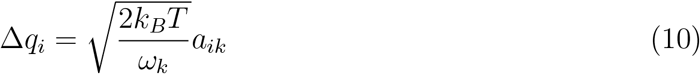

with *a*_*ik*_ the *i*th component of the eigenvector for the *k*th mode of frequency *ω*_*k*_. This expansion was proposed to study proteins and it was implemented for RNA molecules in the present work and it should satisfy Eckart conditions and can be used for computing the overlap and other expressions (see below). See our previous work for the full expressions corresponding to each type of variable in the presence of *N* bodies.^31^

#### Average properties

Thanks to the conversion from ICS to CCS, two average quantities can be computed: the displacements of mean atomic positions from their positions in the minimum-energy conformation, and the mean square fluctuation of each atom from its displaced mean position. As shown in,^31^ if the terms up to second order or more are considered (see eq 9), the conversion from ICS to CCS is no longer linear. The resulting nonlinearity provides a displacement of atomic positions. The average over all conformations is given by

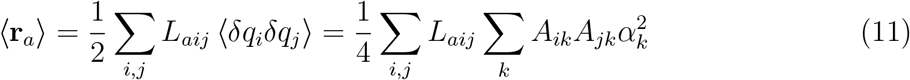

where 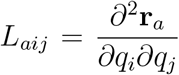 represents the coefficient *ij* (quadratic term) for the **L** matrix. The mean fluctuations are given by

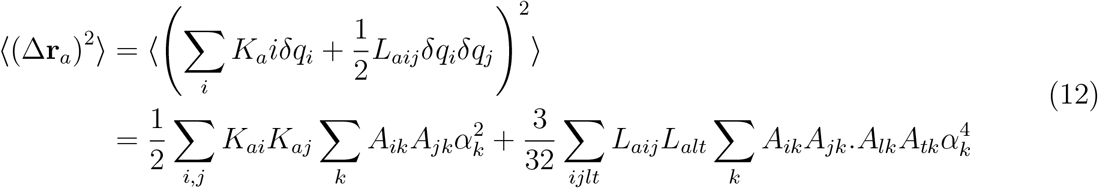

with 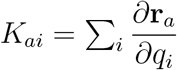.

### Computational details

#### RNA representation

To validate our approach, we used the same coarse-grain (CG) representation proposed in^26^ (hereafter also called RNA three-bead model) to represent RNA molecules. In this model, each nucleotide is described by three pseudoatoms: the first centered on the atom P, the second on the atom C1’ (the ribose) and the third centered on the atom C2 (the nucleobase) (see Figure 1).The mass of each pseudoatom is the sum of the masses of the atoms that constitute it. For the sake of comparison, we also computed iNMA using the coarse-grain model HiRE-RNA proposed by Pasquali and co-workers, where each nucleotide is described by 6 or 7 pseudoatoms (3 atoms for the backbone, 2 for the sugar, 1 or 2 for the nucleobase).^28,58^ For the RNA three-bead model, only the dihedral angles between the pseudoatoms are taken into account (see Figure 1) and each nucleotide has two torsional angles: *τ*_1_ and *τ*_2_. The former involves the torsion around the bond P_*i*_-C1’_*i*_ and the latter is the torsion around the C’1_*i*_-P_*i*+1_ with i the nucleotide under investigation. For the HiRE coarse-grain model, as some of us proposed in^21^ only the torsion angles *τ*_*b,i*_ that involve the backbone are taken into account.

**Figure 1:**
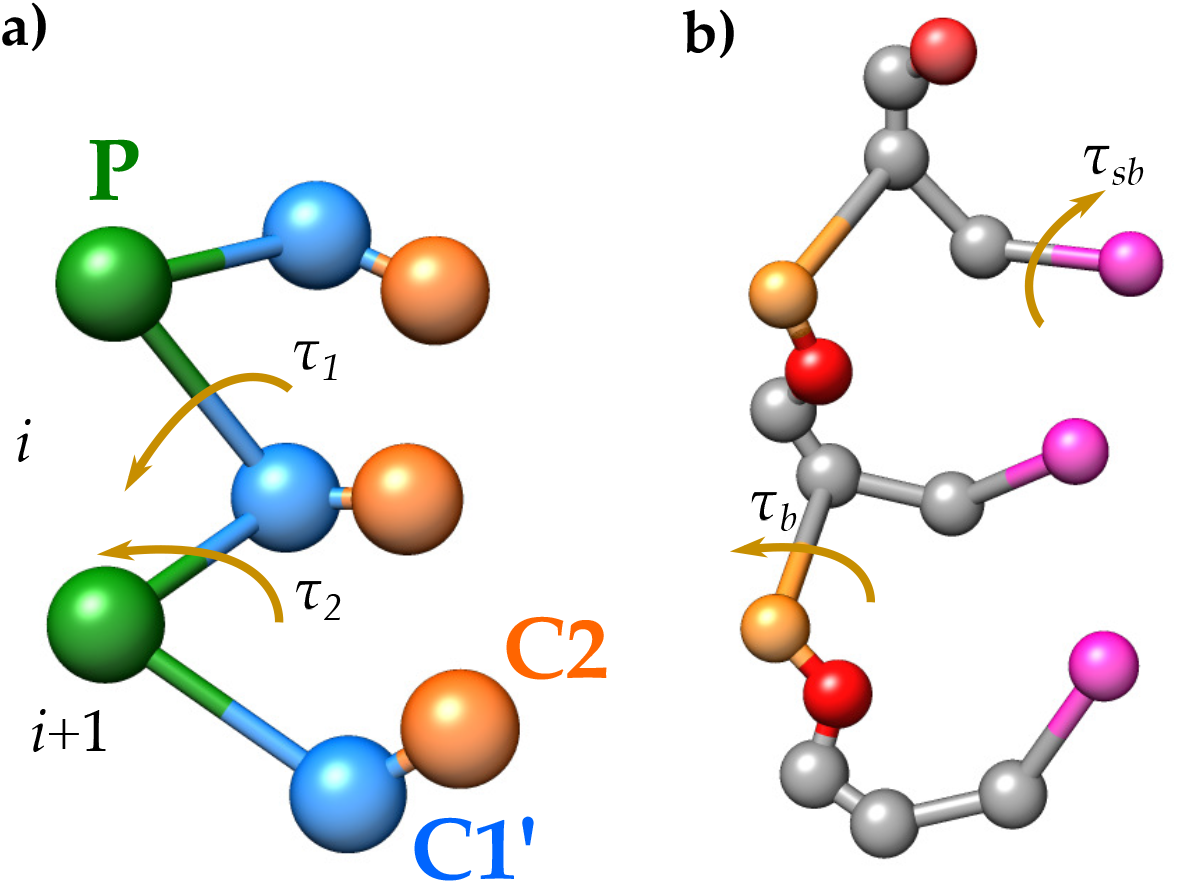
RNA CG representations. a)RNA three-bead model: the green bead represents the pseudoatom centered in P, the blue bead the one centered in C1’ and the orange bead the one centered in C2. The torsional angles *τ*_1_ and *τ*_2_ are also shown. b) HiRE model for three nucleotides. Each bead represents a pseudoatom. An example of the backbone torsion *τ*_*b*_ and the sugar-base torsion *τ*_*sb*_ is shown.

In this work, the HiRE-model was used only for comparison and the full study was conducted using the RNA three-bead model. To avoid artefacts, to compute ICS normal modes we chose the P atom closest to the centre of mass as pivot atom in both models. ICS normal modes were computed using an anisotropic elastic network (ANM) model based on two RNA representations. Two pseudoatoms *i* and *j* are connected if their reference distance 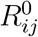 is lower or equal to the distance cutoff *R*_*c*_. Hence, the structure is represented as a network of nodes connected by springs whose force constant is equal to *γ*. The energy of the system was computed based on an ANM defined as follows

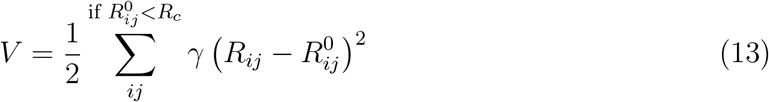

where *R*_*ij*_ represents the distance between the atom *i* and *j*. Here, we investigated the impact of *R*_*c*_ on our results.

#### Calculation Protocol for iNMA

Internal normal mode analysis were computed using different distance cut-offs *R*_*c*_. In each case studied, all the vibrational frequencies obtained were positive (with the exclusion of the six zero-frequency modes representing overall translation and rotation with cNMA calculations). In the case of iNMA, for the first 20 modes within the set { Δ*q*_*i*_} given by eq. (3), we calculated the modified position **r**_*a*_{*q* + Δ*q*} using eq. (9) to perform the conversion from ICS to CCS.

#### All-atom Molecular Dynamics simulations

All-atom MD unbiased simulations were performed for at least 500 ns with the GROMACS 2018 package^59–62^ using the Amber ff99bsc0 force field with state-of-art modification of the force field for RNA molecules.^12^ Each RNA molecule was placed in a dodecahedral box and solvated with TIP4P to a depth of at least 14 Å.^63^ Each system was neutralized by adding potassium cations and then K^+^Cl^−^ ion pairs to reach a physiological salt concentration of 0.15 M and by adding MgCl_2_^64^ based on the experimental conditions. Long-range electrostatic interactions were treated using the particle mesh Ewald method^65,66^ with a real-space cutoff of 10 Å. The hydrogen bond lengths were restrained using P-LINCS,^67^ allowing a time step of 2 fs. Translational movement of the solute was removed every 1000 steps to avoid any kinetic energy build-up. ^68^

Before the MD production, we carried out the energy minimization and equilibration as described in.^17,69^ During equilibration (at least 10 ns) a Berendsen thermostat (*τ*_*T*_ = 1 ps) and Berendsen pressure coupling (*τ*_*P*_ = 1 ps)^70^ were used. The production part was carried out in an NTP ensemble at a temperature held at 310 K and a pressure held constant at 1 bar using the Bussi velocity-rescaling thermostat (*τ*_*T*_ = 1 ps)^71^ and the Parrinello-Rahman barostat (*τ*_*P*_ = 1 ps).^72^ During minimization and heating, RNA heavy atoms remained fixed by using positional restraints. During the equilibration, the restraints were gradually relaxed. Bond lengths were restrained using P-LINCS, allowing a time step of 2 fs. The length of the simulations was at least 500 ns.

#### Principal Component Analysis

In general, the Principal Component Analysis (PCA) is used to reduce the dimensionality on complex data. Here, we applied PCA on both all-atom MD simulations and the ensembles in the Rfam Data Set to get information on RNA dynamics. The first few principal components (PCs) usually capture a significant part of the ensemble variance.

#### PCA on all-atom MD simulations (group 1)

We carried out a PCA in Cartesian Coordinate Space on the all-atom molecular dynamics simulations by considering only the same pseudoatoms of the CG model. To do so, we first superimposed the trajectory converted to CG on the reference structure used for iNMA. PCA was performed by first computing the covariance matrix and then determining the eigenvalues *λ*_*i*_ and the eigenvectors **e**_*i*_ (the principal components) as implemented in GROMACS 2018.^59,60,62^

#### Principal Component Analysis on the RFARM Data Set (group 2)

Also for this data, to reduce the number of important degrees of freedom and simplify the analysis, we conducted a PCA. We constructed a matrix where each row holds all coordinates for a single structure. The structures were superimposed on the reference one. Columns are then variables, one for each structure coordinate. PCA is performed on this matrix using MATLAB 2021a.

### Evaluation normal mode analysis

#### Overlap and Cumulative Overlap

To evaluate the ability of iNMA calculations to correctly predict RNA dynamics, we calculated *O*_*jk*_, the overlap between the conformational change predicted by the *j*th normal mode and the principal component:

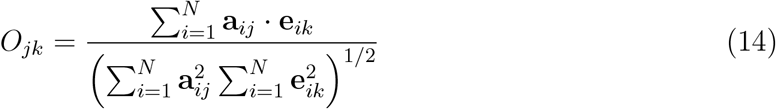

where **a**_**ij**_ represents the *i*th displacement in the *j*-th mode and **e**_*ik*_ the *i*-th displacement in the *k*-th eigenvector of PCA. For iNMA, **a**_**ij**_ is obtained from the ICS to CCS conversion and after the orthogonalization of the normal modes with the Gram-Schmidt process. ^73^

To represent unbound-to-bound conformational changes of proteins that form binary complexes, eq. (14) between the conformational change predicted by the *j*th normal mode and the observed unbound-to-bound conformational becomes:

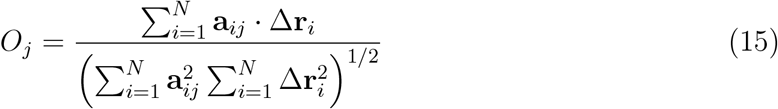

where Δ**r**_*i*_ is the vector describing the unbound-to-bound conformational change for an atom *i*. To compute the vector Δ**r**_*i*_, the unbound and bound structures must be superimposed by weighting the mass following the same approach proposed in.^17,31^

We also computed the cumulative overlap, *CO*_*k*_, to recover the *k*-th principal component *PC*_*k*_ that is defined as

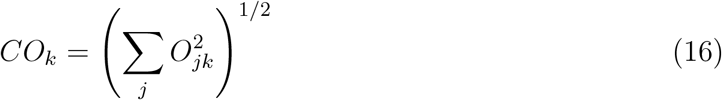

For the unbound-to-bound transition, eq. (16) for the cumulative overlap CO becomes:

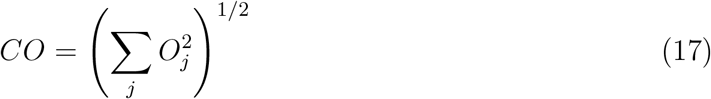

#### Root Mean Square Inner Product

The Root Mean Square Inner Product (RMSIP)^74^ is a measure of the similarity between two sets of eigenvectors (either two sets of normal modes or a set of normal mode and principal components), defined as

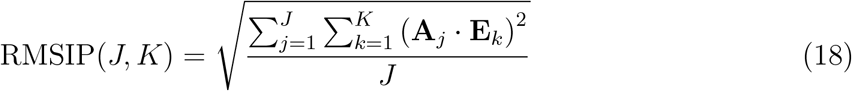

where **A**_*j*_ and **E**_*k*_ are the *j*-th and *k*-th eigenvector obtained by iNMA and PCA, respectively. In other words, it allows to estimate the overlap between the space spanned by the first *I* PCs and the first *J* low-frequency internal modes.

#### Structure Preservation Descriptors

To study large conformational changes, we exploited the possibility of using mode amplitudes beyond those corresponding to room temperature (*α*_*k*_) also for RNA molecules, as proposed for proteins in.^53^ Hence, for the *k* -th mode, the modified amplitude *α*’_*k*_ can be written as

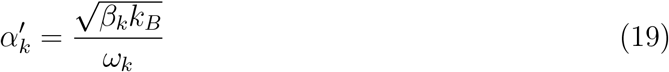

where (*β*_*k*_ is a multiplication factor that is related to the effective temperature, *T*_*eff*_, through the following equation: (*β*_*k*_ = 2*T*_eff_. Another parameter we need to consider is the phase angle, *φ*_*k*_, for a given mode *k* (see eq. (3)). If we consider a single mode, we can set the time equal to zero, and the largest amplitude will be obtained for *φ*_*k*_ = 0° and *φ*_*k*_ = 180°.

Finally, to assess the extent of the structural deformation induced by iNMA modes, we computed (i) the RMSD of the pseudo-atoms in the starting structure and in the structure obtained by the application of normal modes with respect to the target; (ii) the RMSD of the virtual P_*i*_-P_*i*+1_ bonds and 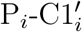 bonds defined as

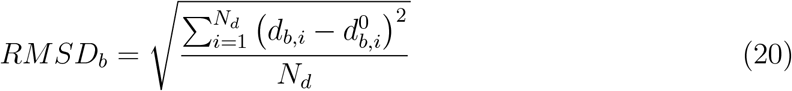

where *b* can either the virtual P_*i*_-P_*i*+1_ bonds or 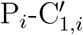 bonds (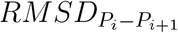 or 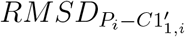), *N*_*d*_ is the number of bonds, 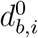 is the reference length of the *i*-th bond *b* in the starting structure and *d*_*b,i*_ is the length of the *i*-th bond *b* in the modified structure.

### Data set

In comparison with proteins, there is a small number of available 3D atomic structures for RNA molecules, as shown in the NDB.^13,14^ Moreover, fewer resources exist for their curation and comparison. In this study, we considered three main data sets: i) one based on single-stranded RNA molecules for which all-atom MD simulations were computed; ii) one based on the available structures belonged to a specific Rfam family; iii) one based on the transition from unbound to bound RNA. The first two datasets was used to validate the capability of iNMA to describe RNA movements and to chose the best parameters for the elastic network model. The third one was used to investigate the capability of iNMA to predict large conformational changes.

#### Single-stranded RNA data set

Based on our previous works^31,53^ and the benchmarks available in the literature for unbound and bound RNA,^75–79^ we selected 16 RNA molecules with different size, function and conformation. For each molecule, first all-atom MD simulations were performed and secondly principal component analysis (PCA) in Cartesian Space were computed. All the details are presented below. Table S1 shows the list of RNA molecules under investigation.

#### Rfam families

In this study, we built our second dataset starting from Rfam^80–83^ an important available resource that collects RNA sequences and available structures into families based on multiple sequence alignments and co-variance models. First, we filtered Rfam families based on the presence or the absence of available 3D atomic structures. For each Rfam family, the number of available 3D structures varies a lot, from few structures up to more than one thousand.

To determine the representative structures for each Rfam family, first we performed a multisequence alignment for all the members whose 3D structure is available using LocARNA.^84–86^ Then, each family is divided in one or more subgroups based on the length and the sequence alignment. Initially, all structures may only differ by 12 bases in length and they must have the same gaps in the alignment. Only groups with 4 or more members are retained. Then the 3D structures are downloaded and the pairwise RMSDs in the initial pool of structures are computed with RNAalign^87^ that is the RNA version of the TM-align algorithm used in the previous study by Zimmermann and Jernigan.^32^ Then, we choose as the seed structure the highest resolution wild-type conformation that is also selected as the first representative. All structures at less than *c* Å RMSD from it are removed from the initial pool. Finally, the structure with the lowest number of neighbors at less than *c* Å RMSD is selected as representative, and all structures at less than *c* Å RMSD from it are removed from the pool. This last step is iterated until the pool of structures is empty. This procedure allows to reduce the structural redundancy within each family: in the set of representative structures, no pair can have less than c Å RMSD. It also allows keeping the maximum number of representatives, since the structure with less neighbors is selected at each iteration. The cutoff *c* is set to 0.5 Å in this study, but we also built representative sets with various *c* values in the [0.1 - 1.5] range, for testing. We chose a cutoff *c* = 0.3 Å for two systems to increase the number of representative structures. The ensemble of structures selected for each Rfarm family differs from the one obtained in.^32^ Table S1 summarizes this second dataset.

#### Unbound-to-bound transition

To build our dataset on unbound-to-bound transition, we used different RNA-protein docking benchmark.^76–79^ Unfortunately, for several complexes, no 3D structure of the unbound RNA is available and the RMSD between the bound and unbound structure is equal to zero, so we also excluded such complexes since the RNA molecule does not undergo conformational changes. Moreover, to do our comparison we need RNA molecules with the same length. Then, we used RNA-align^87^ to compare and superimpose the 3D structures and compute their alignment. Based on all these criteria, we selected 24 RNA molecules whose RMSDs between the unbound and bound structure vary from 2 to 37.1 Å as shown in Table S2.

## Results and Discussion

### Advantages of iNMA for RNA molecules

As we have already shown in our previous works on a large set of proteins and models, iNMA is generally capable of describing large conformational changes and also deals with models.^40^ First, as we demonstrated for proteins, in the same way we computed iNMA on a set of RNA molecules and for a given mode we increased the parameter (*β*_*k*_ to verify if our approach is capable to preserve the structure. To do so, we computed the RMSD of the 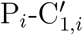 bonds 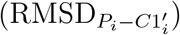 and the virtual P_*i*_-P_*i*+1_ bonds 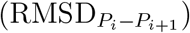 for each RNA molecule. Figure 2 shows an example of our results obtained for the PDB structure 2TRA (A10). As expected, the RMSD computed on the 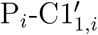 bonds is equal to 0 Å since in our approach the bonds are not taken into account as internal variable. The RMSD for the virtual P_*i*_-P_*i*+1_ bonds slightly increase with (*β*_*k*_ since the structure undergoes a signification conformational change and also it is reasonable that the distance between the phosphates of two consecutive nucleotides may increase.

**Figure 2:**
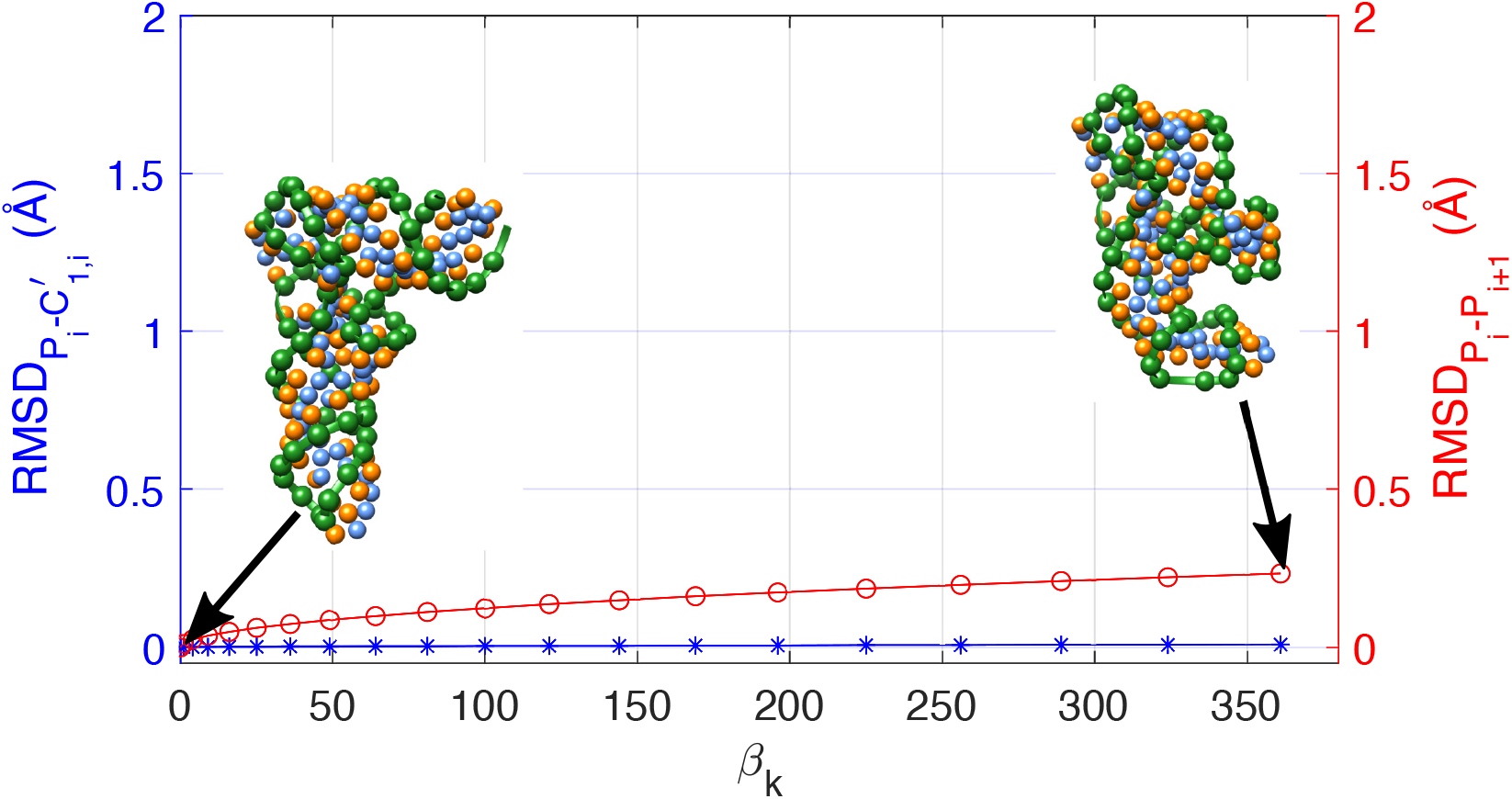
Root-Mean-Square-Deviation (RMSD) for the 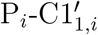 bonds (blue star line)and for the virtual P_*i*_-P_*i*+1_ bonds between two consecutive nucleotides (red dot line) as a function as the multiplicative factor (*β*_*k*_ as defined in eq. (19). The calculations were performed on the PDB structure 2TRA (A10).The starting structure and the final one obtained for the first mode are shown for the sake of clarity.

Second, using iNMA we can also estimate the Root-Mean-Square-Fluctuations (RMSF) per residue using eq. (12). As mentioned before, the RMSF can be computed using a linear or second-order approximation. To evaluate the impact of these approximations, Figure 3 and Figure S1 report the predicted RMSF per residue of the RNA hairpin whose PDB structure is 1QWA and the PDB structure 1L9A, respectively. By comparing Figure 3 and Figure S1 obtained with different values of the force constant *γ*, the larger the fluctuations are large the higher the difference between the approximations highlighting the limitations of the linear approximation for large RMSF. We also investigated the impact of the distance cut-off *R*_*c*_ and the force constant *γ* on the prediction of the RMSF profiles, as shown in Figure 3 for the same RNA structure. Figure S2 shows other examples. We can observe that the RMSF profiles predicted by iNMA show the same flexible regions, but the intensity of the peaks decreases with the increase of the value of the distance cut-off *R*_*c*_. Similar results are obtained if we fix *R*_*c*_ and we change the value of the force constant *γ*. It is important to point out that the distance cut-off *R*_*c*_ not only affects the intensity of the peaks (see Figure 3 B) but also the eigenvectors. On the contrary, the force constant *γ* has an impact mostly only on eigenvalues, and not on the internal modes.

**Figure 3:**
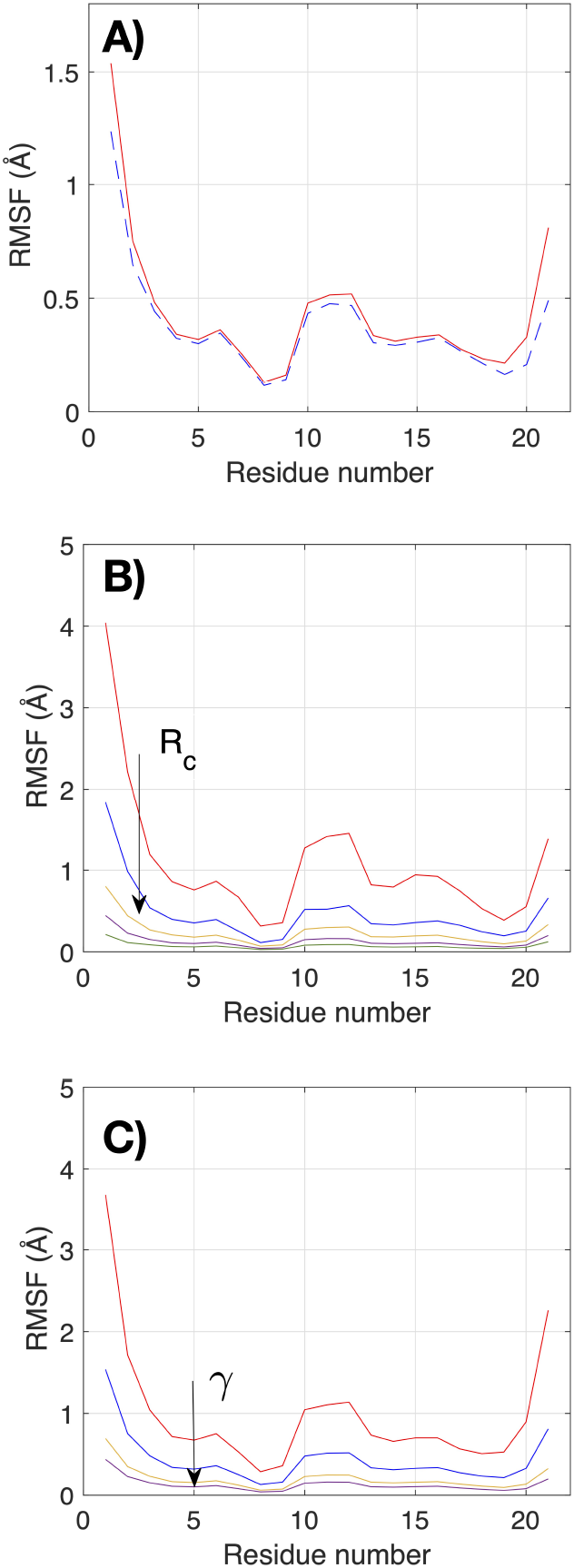
A) RMSF per residue obtained for the PDB structure 1QWA using the first (red dashed line) and second (red) order approximation in eq.(12) and *R*_*c*_ = 16 Å and *γ* = 0.4 kcal mol^−1^. B) RMSF per residue obtained for 1QWA using the following distance cut-off *R*_*c*_: *R*_*c*_ = 10 Å(red),*R*_*c*_ = 12 Å(blue), *R*_*c*_ = 14 Å(yellow), *R*_*c*_ = 16 Å (purple) and *R*_*c*_ = 16 Å (olive green). C) RMSF per residue obtained for 1QWA using the following force constant *γ*: *γ* = 0.1 kcal mol^−1^ (red), *γ* = 0.2 kcal mol^−1^ (blue), *γ* = 0.4 kcal mol^−1^ (yellow) and *γ* = 0.6 kcal mol^−1^ (purple). In the plots B) and C), the arrow indicates the increasing of *R*_*c*_ or *γ* and second-order approximation has been used.

Finally, we can also compare the RMSF per residue predicted by iNMA with that computed from all-atom MD simulations (see Figure 4 for the PDB structure 1QWA and 1L9A, respectively). As highlighted above, based on the distance cut-off *R*_*c*_ and the force constant *γ*, the predicted RMSF profiles change their intensity, although the ratio between the peaks is mostly unchanged. Although we would not expect the iNMA-derived fluctuations to exactly reproduce the all-atom MD results, the peak positions correspond reasonably well. For the PDB structure 1QWA, using a distance cut-off *R*_*c*_ equal to 12 Å and a force constant *γ* equal to 0.6 kcal mol^|1^, multiplying the NMA data by a factor of 4 also leads to a reasonably good agreement in magnitude, with the exception of the ends whose flexibility is overestimated. When a larger value of either the distance cut-off *R*_*c*_ or the force constant *γ* is chosen, a larger multiplicative factor should be used to better fit the magnitude of the peaks as shown in Figure S3. It is important to point out that this good agreement is obtained despite the fact that our analysis is limited to dihedral angles and includes only a simplified energy description.

**Figure 4:**
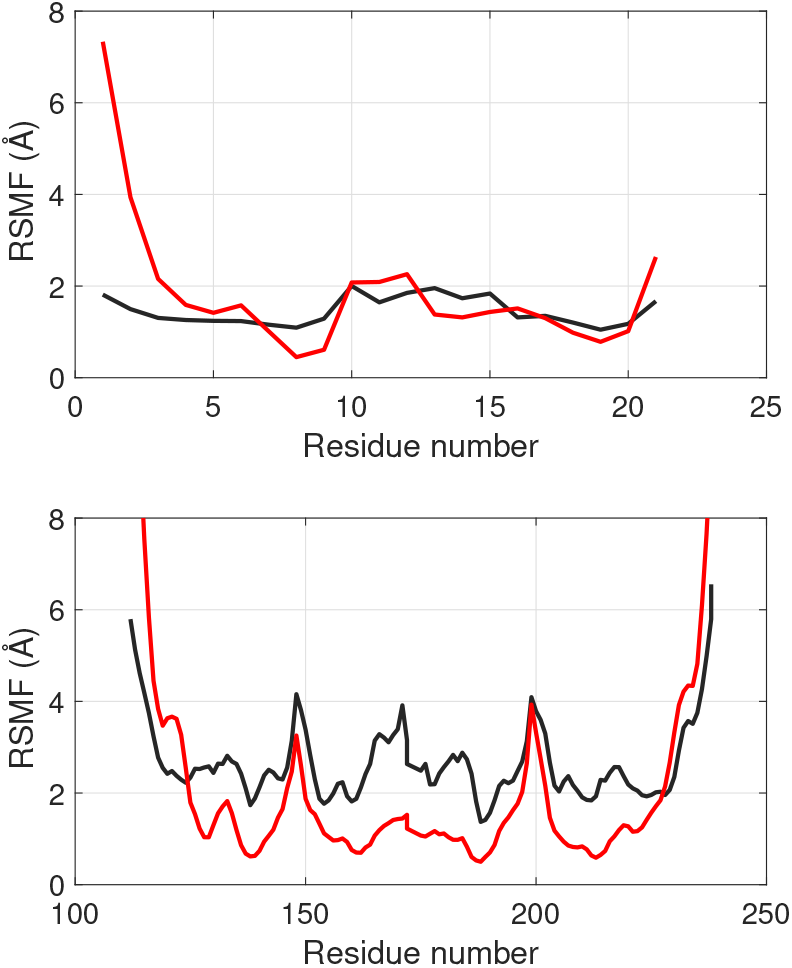
Comparison of iNMA RMSF (red) with RMSF from all-atom MD simulation (black) computed for the PDB structure 1QWA (top) and 1L9A (bottom) using a value of distance cut-off *R*_*c*_ = 12 Å and a force constant *γ* = 0.6 kcal mol^−1^. RMSF profiles obtained by iNMA are multiplied by a factor 4.

As already highlighted in our previous works on iNMA applied to proteins,^17,31,40^ also for RNAs only few residues show significant torsional changes when low-frequency modes are used, but these residues account for a large part of the overall motion. It is important to point out that the knowledge of the key nucleotides responsible for the conformational change is an another intrinsic feature of iNMA, but it is difficult to dermine using Cartesian Normal Mode Analysis. Moreover, thanks to the transformation from ICS to CCS, the location of the largest structural displacements due to the key torsional variables for each mode can be identified. Finally, as already showed we can explore amplitude larger to room temperature without deforming the structure. To highlight the capability of the iNMA to predict large conformational changes, Figure 5 shows the RMSD between the modified structure using internal normal modes and the target structure (RMSD_m-b_) as a function of the multiplicative factor (*β*_*k*_ (see eq. (19)).

**Figure 5:**
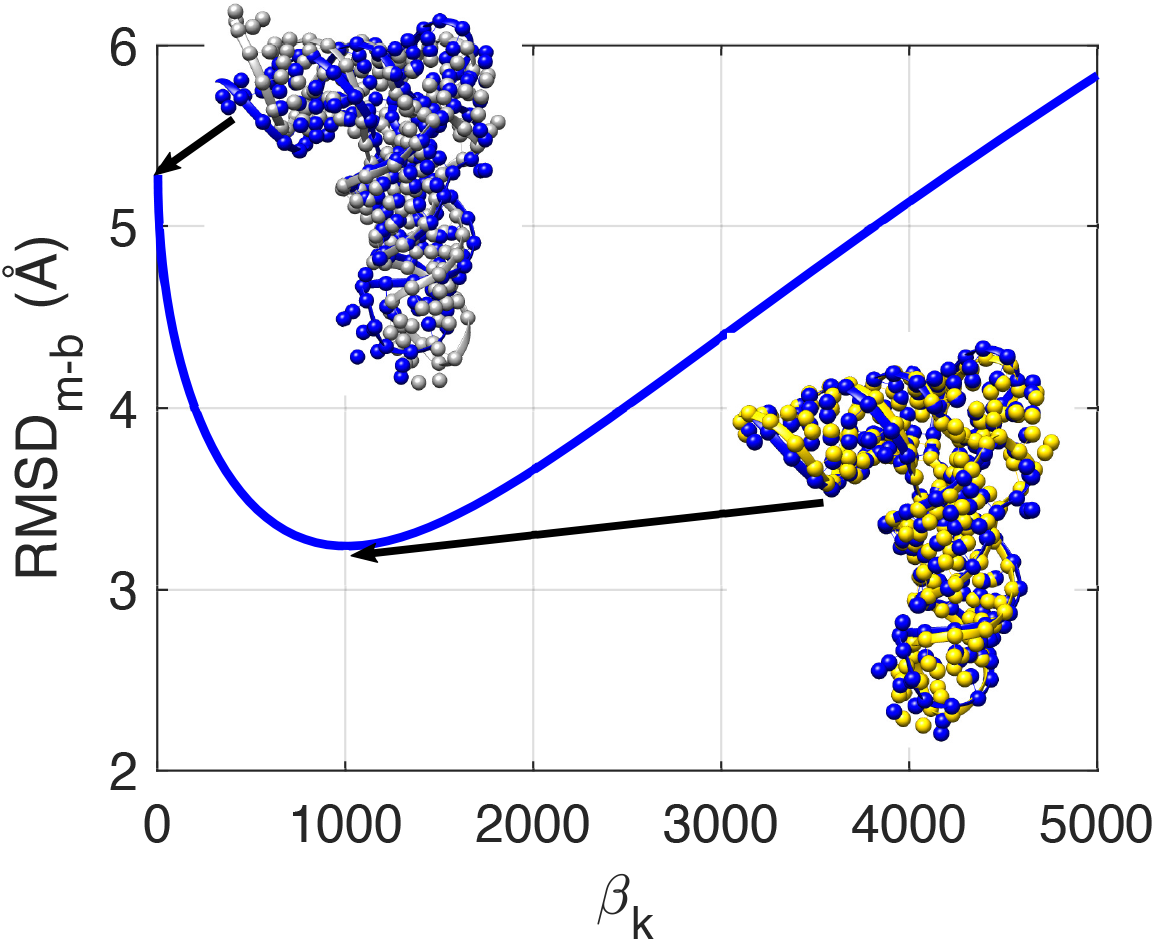
RMSD between the bound structure (PDB ID: 1ASY) and the modified structure after application of mode *j* on the unbound structure 2TRA as a function of the multiplicative factor (*β*_*k*_ (see eq. (19)). In the plot, the unbound structure (colored gray, PDB ID: 2TRA) is superposed to the bound one (colored blue); the modified structure (colored yellow) is superposed to the bound one at the minimum of RMSD.

### Impact of the RNA coarse-grain model on the predictions

Based on the analysis of our results, we chose two RNA molecules to assess the impact of the RNA coarse-grain model on the prediction of flexibility and in particular RNA motions on two opposite scenarios: a compacted structure (a tRNA whose PDB structures is 2TRA) and a completely unfolded RNA molecule like the PDB structure 1SDR. As explained above in the subsection RNA representation, the RNA three-bead model and the model HiRE-RNA were chosen. To compare the normal modes, for both of them we computed the overalp *O*_*jk*_ between the *j*-th internal mode *j* and the *k*-th principal component (PC*k*) obtained by all-atom MD simulations, the cumulative overlap CO_*k*_ and RMSIP. Figure 6 summarizes these results obtained with different value of distance cut-off *R*_*c*_ for the two models, since their resolution is different, we chose *R*_*c*_ = 19 Åfor the RNA three-bead and *R*_*c*_ = 11 Åfor the model HiRE-RNA. For the tRNA, the results are very similar between the two models in terms of cumulative overlap CO_*k*_ and RMSIP, showing that iNMA coupled with the RNA three-bead model and HiRE-RNA can predict 87% and 83% of the motions of the first three dominant PCs by taking in account only the first ten lowest modes. There are just slightly dependent on the values of *R*_*c*_, justifying our choice to use a the three-bead RNA model in this study to investigate quite compact structures.

**Figure 6:**
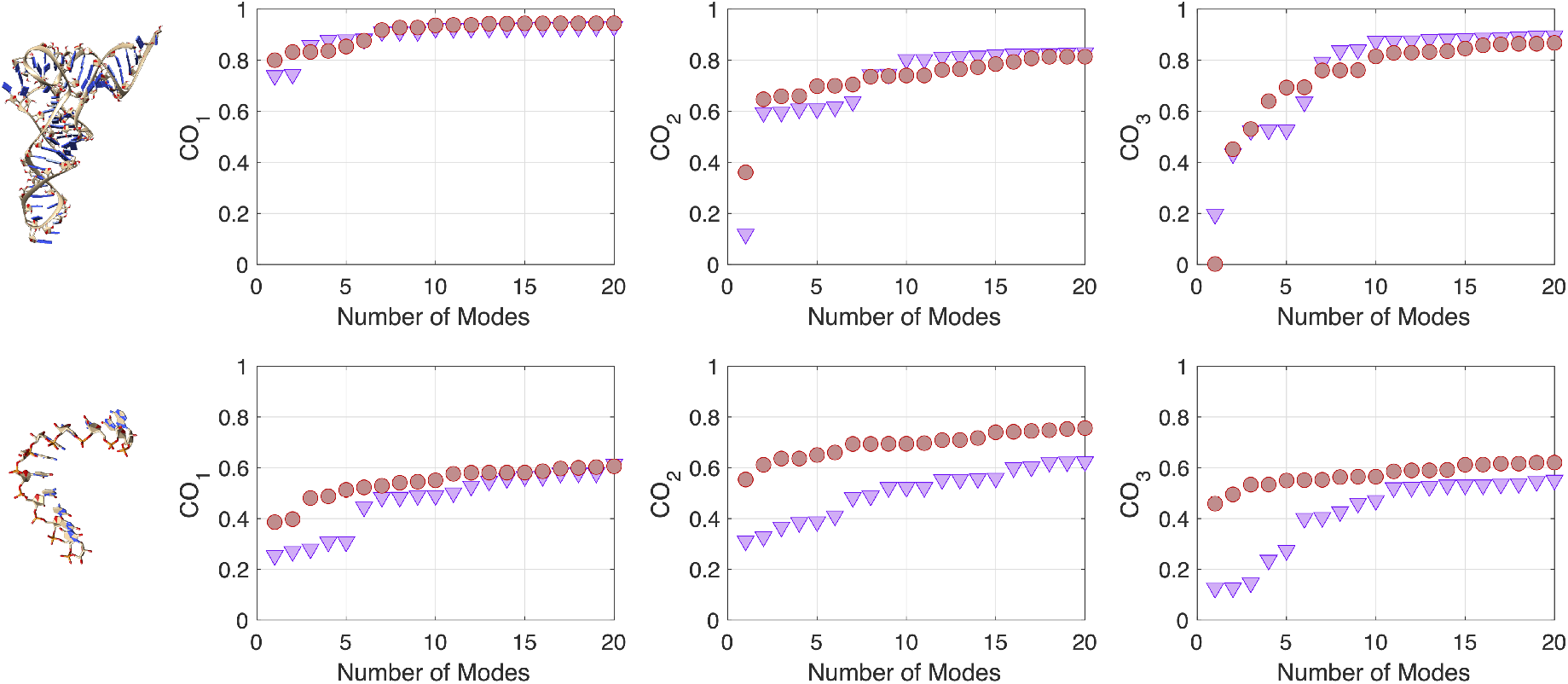
Cumulative overlap (CO) (see eq.(16)) as a function of the number of lowest modes used in the calculation for the first (left), the second (middle) and the third (right) principal components. In top, the results obtained for the PDB structure 2TRA are reported. In the bottom the results for the PDB structure 1SDR. Red circle: HiRE-RNA model. Lilac downward-pointing triangle: RNA three-bead model.*R*_*c*_ = 19 Åfor the RNA three-bead and *R*_*c*_ = 11 Åfor the model HiRE-RNA

Regarding the extended RNA molecule (PDB ID: 1SDR), similar results between the RNA three-bead model and HiRE-RNA are obtained when at least the first six lowest modes are taken into account for the first and third principal component (PC1 and PC3). Although both methods describe at least 50% of motions of the second principal component, the results obtained for HiRE-RNA are a bit better and the difference between the three-bead model and HiRE-RNA becomes constant after the first six lowest modes. A similar trend is observed using RMSIP. By taking into account the first twenty lowest mode, iNMA with HiRE-RNA model can predict 66% of the motions of the first three dominant PCs, and iNMA with the RNA three-bead model 60%. In this case, the results obtained by the three bead RNA model are also impacted by the choice of the value of *R*_*c*_ (see Figure S4 and S5). However, our results are very promising since first predicting the motions of an extended RNA molecule is very challenging and also all-atom MD simulations can suffer of limited sampling. Second, there is a limited lost of information using the three bead-RNA model and a simplified energetic model, mostly due the intrinsic advantages of iNMA.

### Comparison of predicted flexibility obtained by iNMA with structure ensembles

#### MD ensembles

First, to assess the capability of internal modes computed by iNMA to properly described the flexibility and the motions observed in all-atom MD simulations (here-by termed MD ensembles), we studied the first dataset (dataset 1) and for each system, we computed the overlap *O*_*jk*_ and the cumulative overlap CO_*k*_ for value of the distance cut-off *R*_*c*_ ranging from 11 Å to 16 Å. Figure S15 shows an example of the matrix of overlaps *O*_*jk*_ obtained for the PDB structure 2TRA. Interestingly, for some systems, only a mode seems sufficient to recover completely a specific principal component (PC). In particular, the maximum overlap *O*_*j*_ between the first low-frequency internal mode and the first principal component show a range of values of 0.16 → 0.94. For PC2 and the lowest mode, this range of values varies between 0.19 and 0.92 and for PC2 and the second lowest mode between 0.12 and 0.82. Figure S16 represents the maximum overlap *O*_*jk*_ obtained for the first three PCs and Figure 7 shows the location of the maximum (i.e. the best) overlap *O*_*j*1_ between the internal modes and PC1 in the lowest twenty mode. The iNMA approach places the maximum overlap between the internal mode and the first principal component within the first lowest of mode in ∼ 44% of the cases, within the two lowest modes in ∼ 69% of the cases and within the five lowest mode in ∼ 81% of the cases suggesting that a few modes are sufficient to properly describe the motions.

**Figure 7:**
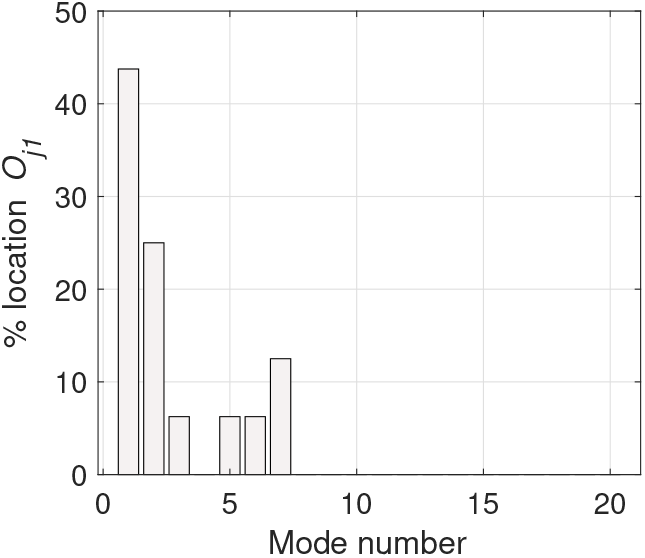
Percentage location of the maximum overlap, *O*_*j*1_, obtained for the first principal component PC1 on dataset 1, for the 20 lowest frequency modes.

Figure 8 illustrates how the cumulative overlap changes as a function of the value of the distance cut-off *R*_*c*_, the number of modes taken in account (the first ten and 20 low-frequency internal modes) and the system under investigation for the first principal component (PC1). Figure S17-S21 show the results obtained for PC1 using the five, ten and fifteen lowest-frequency internal modes, for PC2 using the first five, ten, fifteen and twenty low-frequency modes and for PC3 using the first five, ten, fifteen and twenty low-frequency modes. For some molecules of RNAs, COs can be affected by the choice of the value of the force constant *R*_*c*_ showing a complicated trend. By considering the lowest ten modes, we can predict between 43% and 93% of the motions of the PC1 by choosing *R*_*c*_ = 16 Å. Again, these results suggested that few modes are necessary since the cumulative overlap slightly improve if the first twenty low-frequency modes are considered instead of the first ten ones.

**Figure 8:**
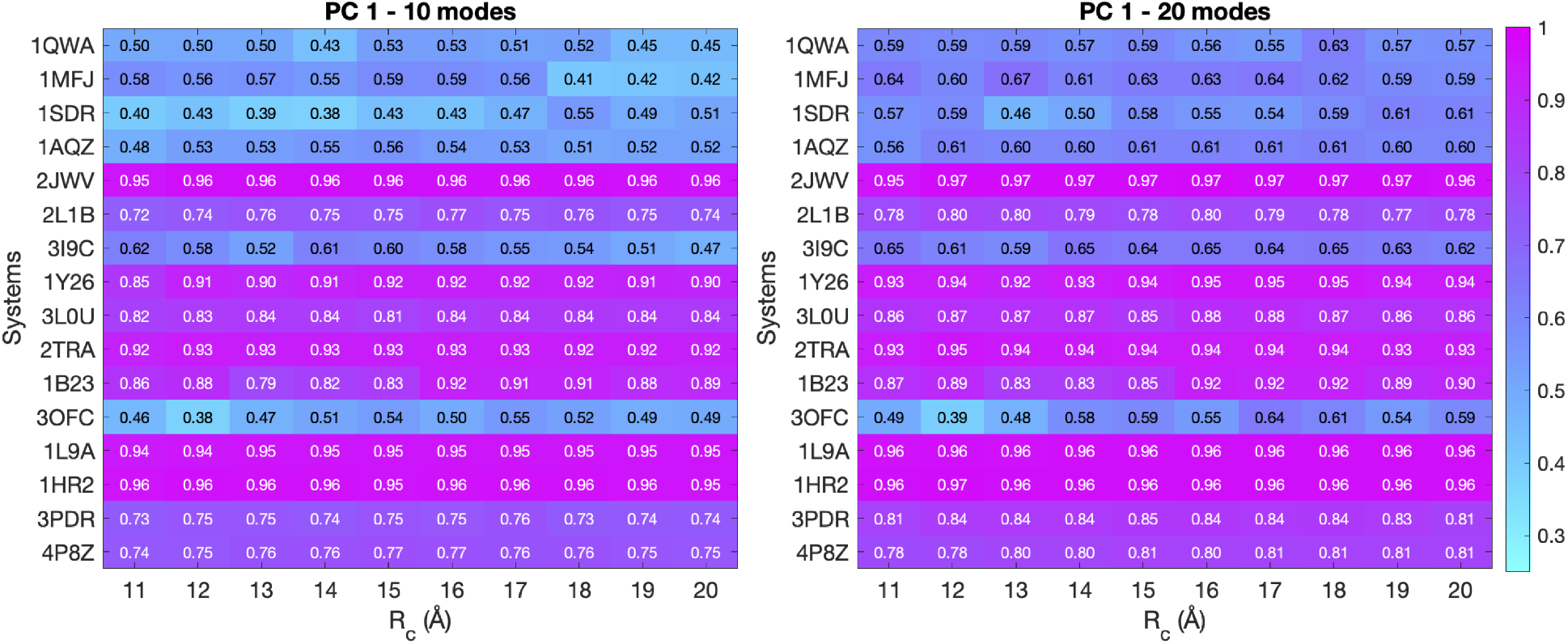
Cumulative overlap (CO) (see eq. (16)) computed using the first ten (left) and twenty (right) low-frequency modes and the first principal component (PC1) for all the systems under investigation in dataset 1. The color of each cell is based on the calculated CO and goes from cyan (low CO, meaning that less than 30% of the motions are captured) to magenta (CO equal to 1, i.e., all the motions are captured and recovered).

To better characterize these results, we chose a cut-off of 0.5 for the cumulative overlap computed on the first principal component (PC1). By setting this cut-off, the percentage of systems that capture at least 50% of the dominant motions varies with the value of the distance cut-off *R*_*c*_ and the number of modes taken into account, as summarized in Figure 9. The percentage rapidly increases until 10 modes where around 94% of systems captures at least 50% of the dominant motions using a distance cut-off equal to 16 Å. A similar trend is observed also for both PC2 and PC3 (see Figure S13). In case of PC2, using the lowest ten internal modes for all the systems we can describe at least 50% of motions. Although using ten modes, the best results are obtained using a distance cut-off equal to 14 Å, similar results are also obtained with *R*_*c*_ equal to 16 Å. Finally, similar results are obtained by analysing the comparing iNMA and the combination of the first three PCs via RMSIP (see eq.(18)). Using only the first five low-frequency modes, iNMA can capture between 31% and 100% of three dominant PCs (PC1, PC2 and PC3). Figure S23 summarizes these results by showing the percentage of systems which captures at least 50%, 60%, 70% and 80% of first three dominant PCs.

**Figure 9:**
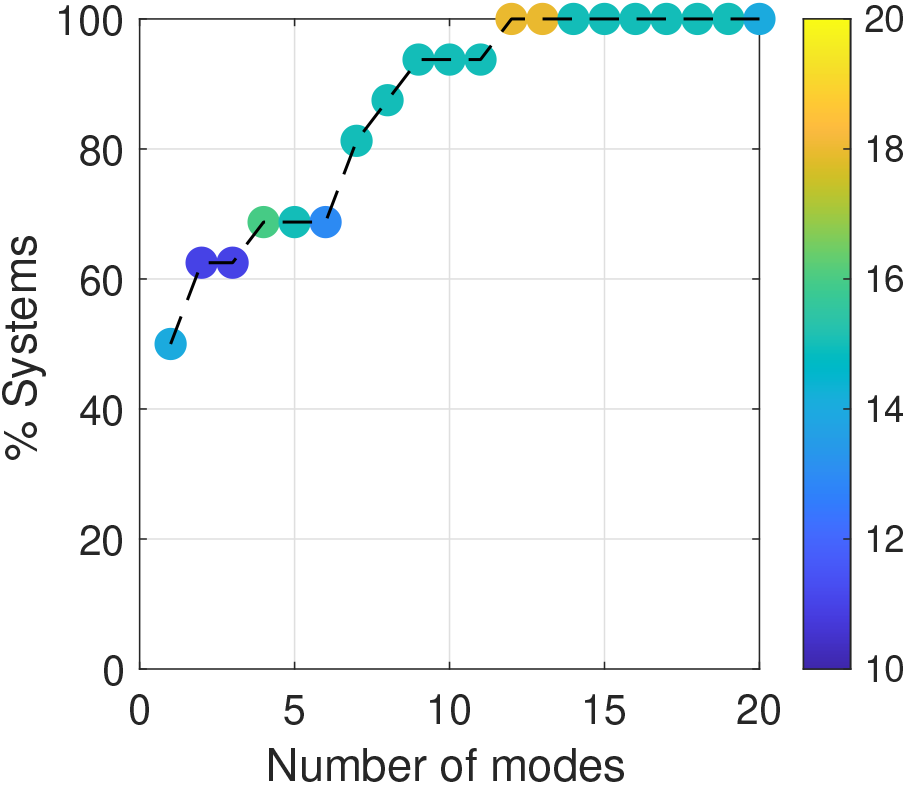
Percentage of systems of dataset 1 which capture at least 50% of the motions of the first principal component as a function of the number of modes taken into account. The dots are colored based on the value of the distance cut-off *R*_*c*_ from blue to yellow for which the best prediction is obtained, for example, *R*_*c*_ = 16 Å for green dots.

#### Experimental ensembles from Rfam families

Using the same strategy, we also compared the predicted flexibility obtained by iNMA with those described by experimental structure ensembles (dataset 2 in Table S1). To do so, we considered the second dataset based on the Rfam families for which 3D structures are available. For each system, we performed iNMA on the representative structure for value of the distance cut-off *R*_*c*_ ranging from 11 Åto 16 Åand we computed the overlap *O*_*jk*_ and the cumulative overlap CO_*k*_. Figure S6 shows an example of the matrix of overlaps *O*_*jk*_. As for the MD ensemble, for fewer cases only a mode seems sufficient to recover completely a specific principal component (PC). In particular, the maximum overlap *O*_*j*_ between the first low-frequency internal mode and the first principal component show a range of values of 0.05 → 0.64. This range of value is lower than the one obtained for MD ensembles, and we can speculate that this is due to the lack of experimental structures in some ensembles or the overrepresentation of some conformations. For PC2 and the lowest mode, this range of values varies between 0.05 and 0.75 and for PC2 and the second lowest mode between 0.02 and 0.53. These results are similar to the ones obtained for MD ensembles. Figure S7 summarizes these results by showing the maximum overlap *O*_*jk*_ obtained for the first three PCs. In comparison with the work of Zimmermann and Jernigan^32^ conducted on a similar dataset where 3 modes are necessary to describe the CO for their case study (the tRNA family, here system R1), our findings highlight again the potential of iNMA to predict RNA flexibility and motions. In fact, the iNMA approach places the maximum overlap, *O*_*j*1_, between the internal mode and the first principal component within the two lowest modes in ∼ 48% of the case and within the five lowest mode ∼ 81% (Figure 10).

**Figure 10:**
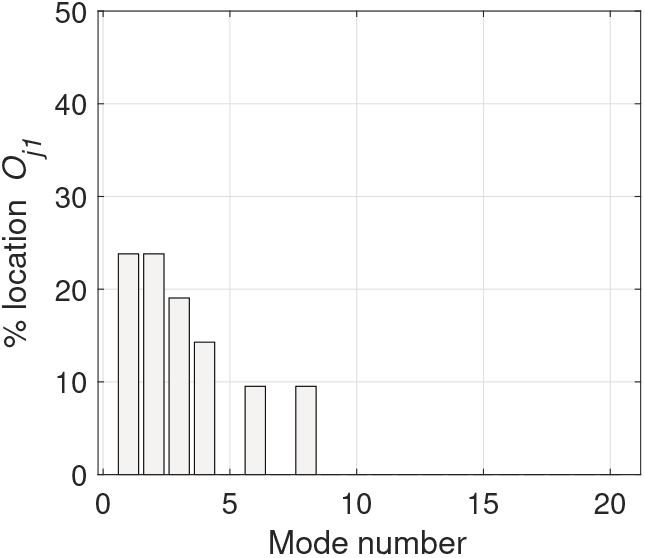
Percentage location of the maximum overlap, *O*_*j*1_, obtained for the first principal component PC1 on dataset 2, for the 20 lowest frequency modes.

Figure 11 illustrates how the cumulative overlap changes as a function of the value of the distance cut-off *R*_*c*_, the number of modes taken into account (the first ten and 20 low-frequency internal modes) and the system under investigation for the first principal component (PC1). Figures S8-S12 show the results obtained for PC1 using the five ten and fifteen low-frequency internal modes, for PC2 using the first five, ten, fifteen and twenty low-frequency modes and for PC3 using the first five, ten, fifteen and twenty low-frequency modes. A slight dependence on the choice of the distance cut-off is observed, as obtained in previous analyses on MD ensembles. To assess the capability of the iNMA approach to predict experimentally observed RNA motions, we also compared our results with the ones presented in,^32^ although we redefined the ensembles using our new approach. In both studies, the first sixteen families are the same. First, we can observe that our approach has similar results and for several families (∼ 75% of ensembles in common) our predictions are better than the ones reported by Zimmermann and Jernigan in Figure 1 in.^32^ However, for the family of 5.8sRNA, our predictions are less good than theirs, but this discrepancy can be explained by the fact that our new ensemble differs from the one used in.^32^ Based on our refinement approach, we almost halved the number of structures for several families with respect to,^32^ and we can suggest that the differences may arise from the lack of 3D structures in the new ensemble. However, based on the overall results and the advantages of iNMA already presented, our approach seems more suitable than cNMA to study the dynamics of RNA ensembles.

**Figure 11:**
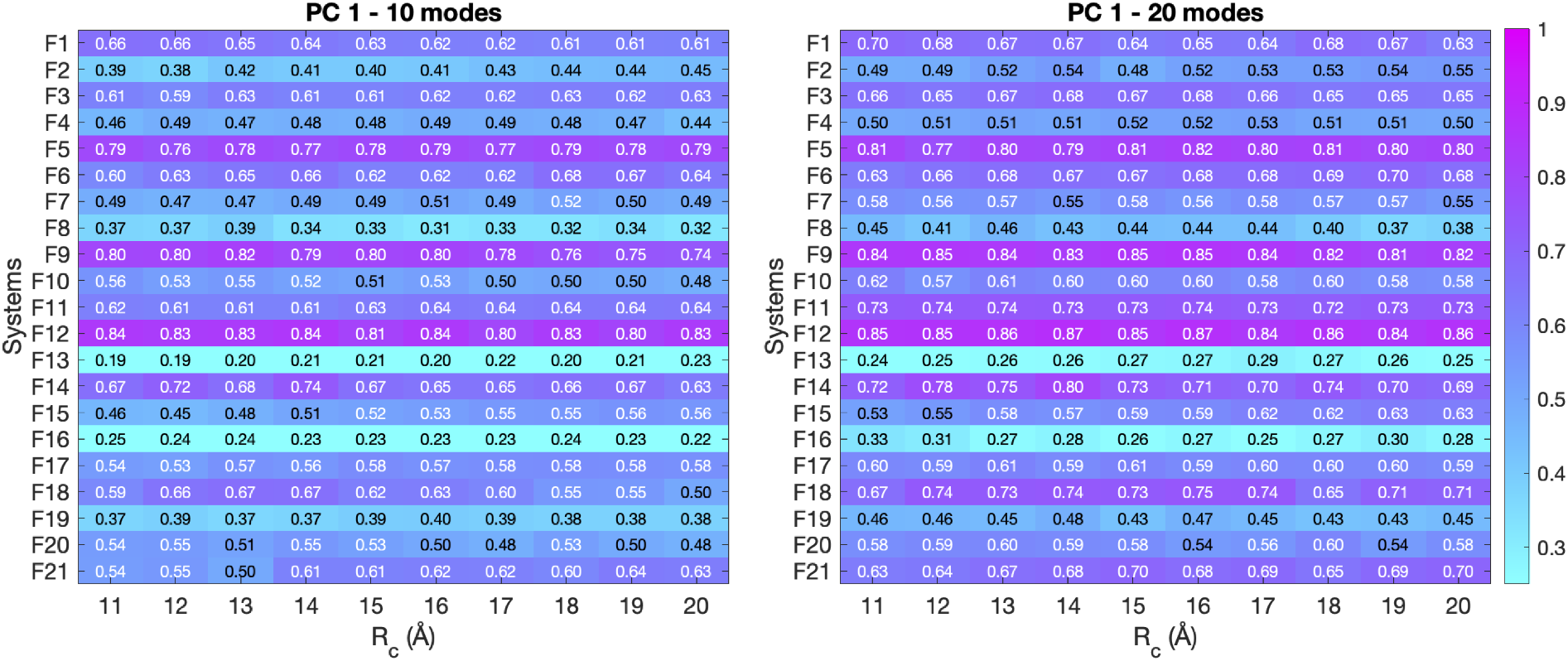
Cumulative overlap (CO) (see eq. (16)) computed using the first ten (left) and twenty (right) low-frequency modes and the first principal component (PC1) for all the systems under investigation in dataset 2. The color of each cell is based on the calculated CO and goes from cyan (low CO, meaning that less than 30% of the motions are captured) to magenta (CO equal to 1, i.e., all the motions are captured and recovered).

As for the ensemble obtained by MD simulations, to better characterize these results, we chose a cut-off of 0.5 for the cumulative overlap computed on the first principal component (PC1). By setting this cut-off, the percentage of systems that capture at least 50% of the dominant motions varies with the value of the distance cut-off *R*_*c*_ and the number of modes taken into account, as summarized in Figure 12. The percentage rapidly increases until 10 modes where 72% of systems captures at least 50% of the dominant motions using a distance cut-off equal to 16 Å. A similar trend is observed also for PC2 (see Figure S13). Although for 10 modes, the best results are obtained using a distance cut-off equal to 11 Å, similar results are also obtained with *R*_*c*_ equal to 16 Å. For PC3, our results highlight the need to use up to 20 modes to recover the motions (see Figure S13), when on the contrary it is not necessary for MD ensembles. To understand this discrepancy, we can make two hypothesis: i) although the third PCs is dominant, it is already more sensible to small differences as also observed when the maximum overlap *O*_*jk*_ is computed hence the need of a large number of internal normal modes to recover this motion; ii) the lack of structures can also influenced our capability to predict RNA motions. Finally, we also compare iNMA and the combination of the first three PCs via RMSIP (see eq. (18)) and results are are similar. In fact, iNMA can capture between 26% and 68% for the first five low-frequency modes, between 32% and 72% for the first ten low-frequency modes and between 40% and 80% for the first twenty low-frequency modes of three dominant PCs (see Figure S14 for the percentage of systems which captures at least 50% of three dominant PCs).

**Figure 12:**
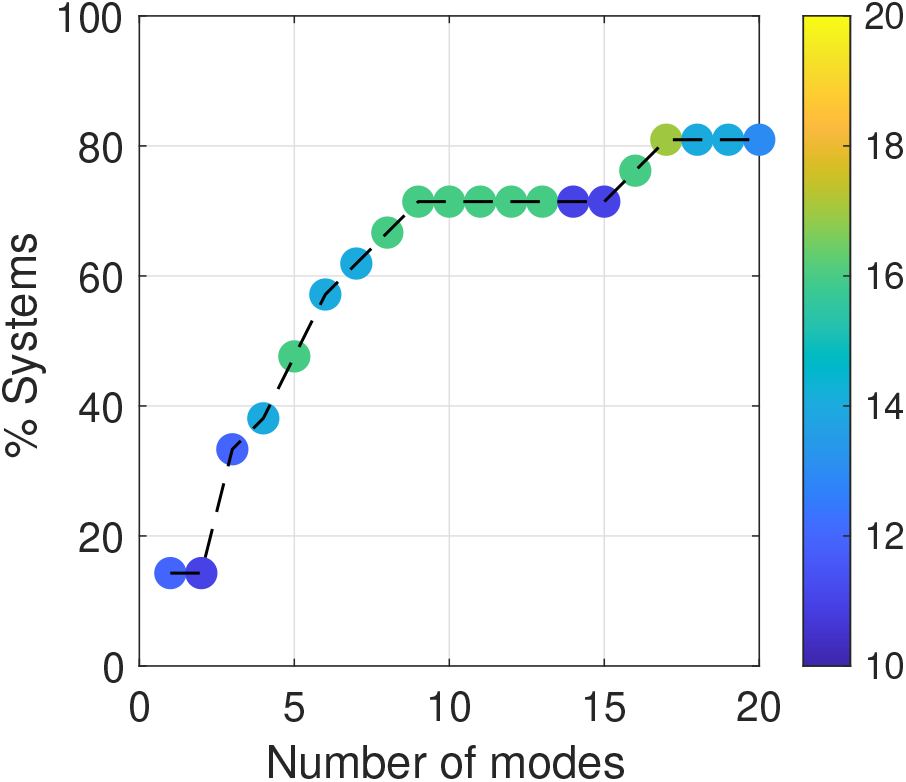
Percentage of systems of the dataset 2 which capture at least 50% of the motions of the first principal component as a function of the number of modes taken into account. The dots are colored based on the value of the distance cut-off *R*_*c*_ from blue to yellow, For example, *R*_*c*_ = 16 Å for green dots.

### Study of unbound-to-bound transition

In order to study the unbound-to-bound transition, based on the results presented above, we chose a value of 16 Å for the distance cut-off *R*_*c*_ to perform iNMA of the RNA molecules in dataset 3 (see Table S2). First, we evaluated the overlap *O*_*j*_ and the cumulative overlap, CO. Figure 13 summarizes the maximum, *O*_*j*_, and the cumulative overlap, CO, as a function of RMSD for the lowest 20 modes of a full set of 24 RNA molecules that we have studied using iNMA. Considering first the maximum overlap, we observe a range of values of 0.14 → 0.73, that is close to the one obtained in our previous work for the proteins.^31^ It is important to point out that the iNMA approach continues to provide good results, even for RNAs that undergo large conformational changes. Moreover, since the normal modes represent an orthonormal space, we can easily combined them and compute the cumulative overlap, as shown above. Once again, we observe that iNMA is a suitable method to describe the transition from unbound to bound structures and, for some RNAs, we reach a cumulative overlap equal to 1 (i.e., a complete description of the transition unbound-to-bound conformation) including for RNAs undergoing large conformational changes. Figure 13 also shows the average cumulative overlap, ⟨CO⟩_*j*_, as a function of the *j* number of modes used in the calculation. As expected, the cumulative overlap increasing by adding the modes and with 10 modes it already reaches an average value of 0.6. Based on the plot the first 5-10 modes provides the larger contributions, suggesting that only few modes are needed as shown before to describe the flexibility.

**Figure 13:**
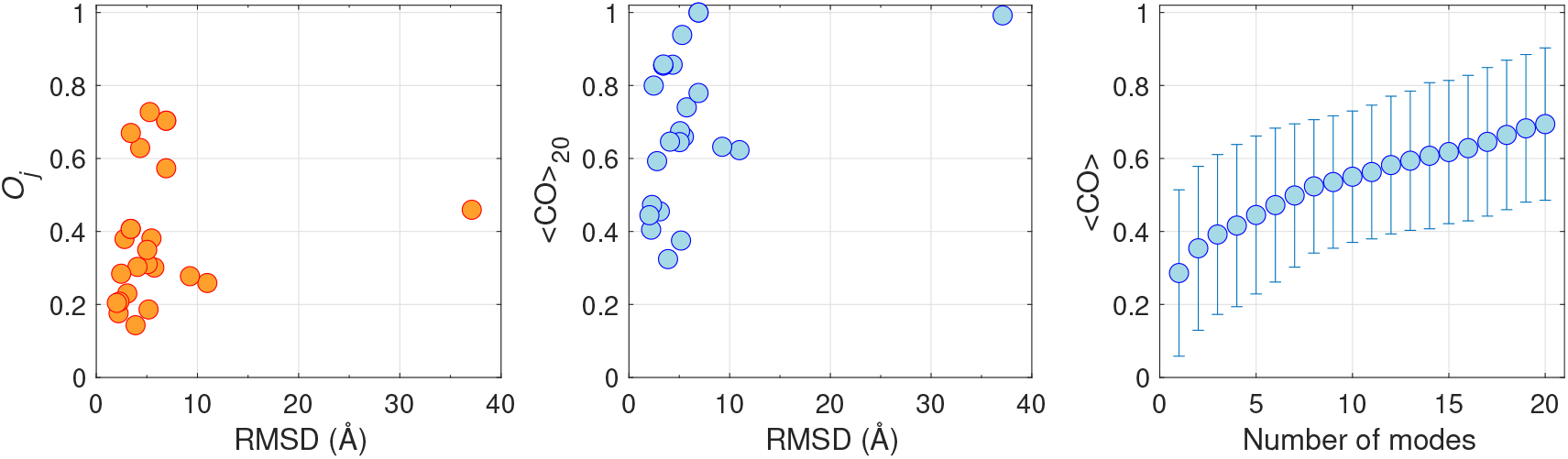
Maximum overlap, *O*_*j*_ (left) and cumulative overlap *CO* (middle) for 24 RNAs proteins, as a function of the RMSD between the unbound and bound structure, computed using the 20 lowest modes obtained. Right: Average cumulative overlap ⟨CO⟩ computing with increasing number of internal normal mode. The error bar indicates the standard deviation of these values.

As already shown for proteins,^17,31^ these results obtained for RNA molecules confirm that unbound RNA structures can often be perturbed along a single low-frequency mode to produce a conformation that is close to the bound one. This behavior can advantageously be used in docking algorithms to account for RNA flexibility in an efficient way or in integrated approaches as internal variables. However, two important problems remain: i) *how should we chose the “correct” mode(s) and how many?* ; ii) *how closely the target conformation can be reached using iNMA?* To answer the first questions, the overlap and the RMSD between the modified structure by internal normal modes and the bound structure can be taken into account. Figure 14 illustrates the location of the best mode in term of overlap *O*_*j*_, and the cumulative results for the first 20 modes. As for proteins,^31^ also for RNAs, the iNMA approach places the maximum overlap within the five lowest modes in ∼67% of the cases, and, moreover, shows a monotonically increasing probability of finding the maximum overlap as the frequency decreases within the first five modes.

**Figure 14:**
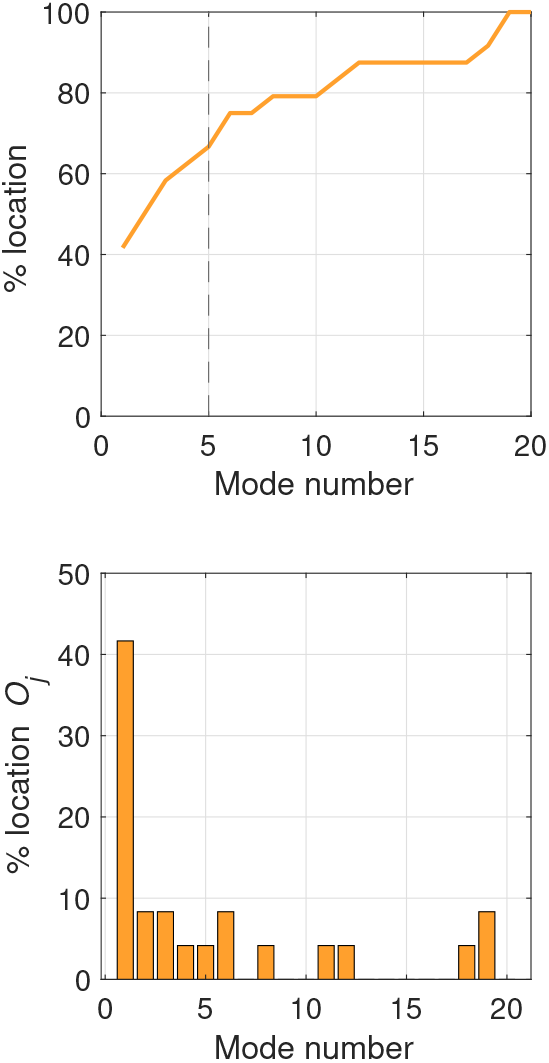
Percentage location of the maximum overlap, *O*_*j*_, (top) and percentage location of the maximum overlap, *O*_*j*_ (bottom) obtained on the dataset of unbound-to-bound transition (dataset 3, Table S3), for the 20 lowest frequency modes.

However, the prediction of the best mode based on the overlap does not guarantee that the selected mode is still the best one in terms of RMSD, when amplitudes of movement above room temperature are considered. Hence, first, for each mode we also computed the modified structures using different multiplicative factor (*β*_*k*_ ranging from ∼ 70 to ∼ 60, 000 and then we determined the best mode by computing the optimal RMSD between them and the target structure (see Table S3 for a summary). The results obtained using overlap (Figure 14) or RMSD (Figure 15) naturally differ, but, in fact, follow similar trends. In particular for RMSD, studying the first two modes is sufficient in many cases (63%). Taking into account the first five modes increases the value to ∼ 75%.

**Figure 15:**
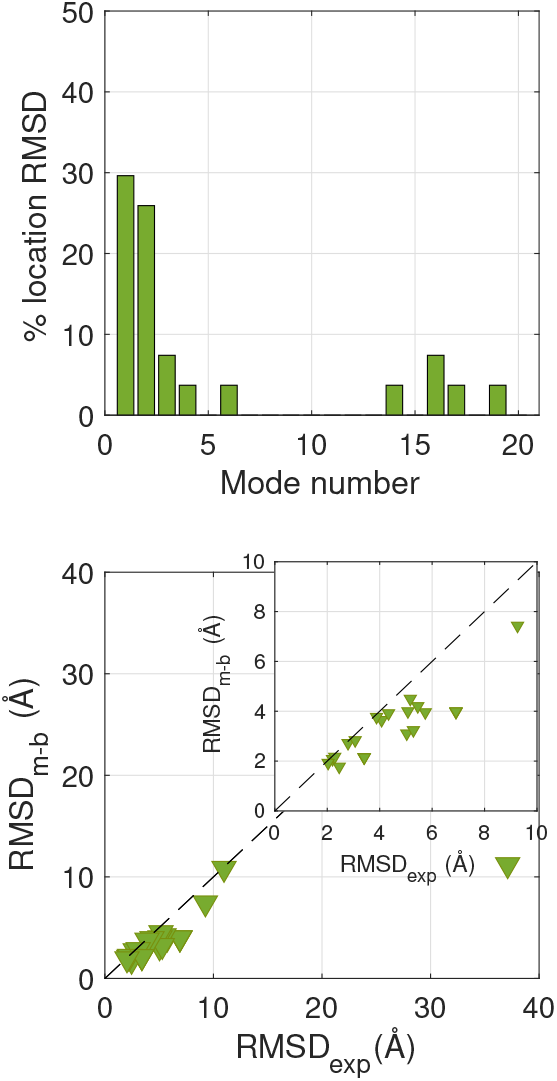
Top: Percentage location of the best mode for RNA molecules in dataset 3 (Table S2) based on RMSD at optimal multiplicative factor (*β*_*k*_. Bottom: Optimal RMSD between the modified structure using the best single iNMA mode and the target (RMSD_m-b_) plotted versus the RMSD between the unbound and bound structures (RMSD_exp_). The dashed line represent the case where no change in RMSD is obtained after application of an internal mode (RMSD_*m−b*_=RMSD_exp_). The insert plot is a zoom for the RNA molecules whose RMSD_exp_ is between 0 and 10 Å.

These results suggest that the best mode describing a transition between two conformations is likely to be amongst the first five and, often, the first two, however we do not answer the second question since they cannot quantify how closely we can reach the target using iNMA. To address this point, first we can determine the *RMSD*_*m-b*_, the RMSD of the modified structure obtained using the best internal normal mode among the first twenty ones with the target at the optimal value of the multiplicative factor (*β*_*k*_ and second compare RMSD_*m-b*_ versus RMSD_*exp*_ the experimental RMSD between the experimental starting (i.e., unbound) and target structures (i.e., bound), as shown in Figure 15. To evaluate our results, we can also compute the percentage of RMSD change defined as 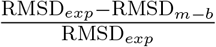. By setting a cut-off of 20% for Δ RMSD, 54% of RNAs have modified structures that are within this varation of RMSD distance of the target, when the best mode is chosen. This value slightly varies until a cut-off of 40% for Δ RMSD, where only four RNAs (17 %) undergo a change in the RMSD with respect to the bound structure of 40%. However, these results are very encouraging as the conformation changes we are trying to predict cover a range up to 40 Å and only a mode is taken into account, as proof of concept.

## Conclusions

In this work, we have described the implementation of normal mode analysis in internal coordinate space (iNMA) for RNA molecules coupled with a simple description of RNA topology (the RNA three-bead model) and a simplified energetic potential. Our implementation is based on our previous works on protein and protein complexes, and it presents several advantages. Among them, a notable advantage is that, even for large movements, internal normal modes avoid the damage to bond lengths and valence angles that occurs with Cartesian coordinate normal modes.

We have applied iNMA to predict RNA flexibility and unbound-to-bound transitions. To do so, we set up three separate datasets. The first one includes 15 all-atom MD simulations of RNA molecules, the second one contains 21 ensembles of experimental 3D structures for different RNA families and the third one includes 24 RNA molecules for which the unbound and bounded 3D structure is known. The first two datasets allowed us to compare the predicted flexibility by iNMA and the one obtained by MD simulations and in experimental ensembles. The third dataset aims to study the unbound-to-bound transition for RNA molecules.

To assess the capability of iNMA to well capture RNA flexibility we computed several quantities: the overlap *O*_*jk*_ per internal mode *j* and principal component *k*, the cumulative overlap, CO_*k*_, for the 20 lowest-frequency modes with respect to the *k*-th principal component and the Root Mean Square Inner Product (RMSIP) between the first 3 principal components and the 20 lowest-frequency modes. First, the most relevant modes are found within the five lowest frequency modes in ∼ 80% of the cases studied. Second, our results are slightly affected by the value of the distance cut-off *R*_*c*_. The approach iNMA using the 20 lowest modes is capable to describe at least 50% of the motion of PC1 for almost all RNA molecules and 80% of them in dataset 1 and dataset 2, respectively. The discrepancy between dataset 1 and dataset 2 may be due to the lack of some structures in some experimental ensembles or the overrepresentation for some conformations. Interestingly, these results are slightly affected if the ten lowest modes are taken in account suggesting that few modes are sufficient to describe the flexibility. For two RNA molecules, we also computed iNMA using a more sophisticate coarse-grain represent (HiRE-RNA). By comparing the results, for a compact structure no significant differences are obtained. For a completed unfolded structure, the differences are observed only if few modes are taken into account suggesting that the use of three-bead RNA model with iNMA allows to retain a large part of structural information and iNMA is a more suitable approach than cNMA to study loosely packed structures.

To analyse the unbound-to-bound transitions, we also computed the overalp *O*_*j*_, the cumulative overalp CO and the RMSD between the modified structure obtaiend after application of the mode *j* and a multiplicative factor (*β*_*k*_ and the bound experimental structures. The iNMA approach seems to require few modes to represent the conformational transitions. Again, the most relevant modes are found within the five lowest frequency modes in ∼ 70% of the cases studied by considering both the overlap O_*j*_ and the RMSD. Moreover, the probability of finding the single optimal mode increases monotonically with decreasing frequency. The cumulative overlap of the the lowest twenty modes can also reach unity. The iNMA approach can predict large conformational changes also for RNA molecules like the ones are expected to occur for RNAs that binds the ribosome.

These promising results open a new route to treating RNA flexibility for unbound structures as well during the formation of RNA-ligand/protein interactions. By adding only few internal modes, it should be possible to implementing iNMA in the refinement of structures in combination with data from low-resolution techniques such as SAXS, in integrative approaches to assembling large molecular complexes and in docking methodologies.

## Author Contributions

The study was designed by EF. Internal Normal Mode calculations were performed and analyzed by TGCB, RS and EF. RS, JM and EF created the three dataset. AS and EF performed all-atom MD simulations and PCA. AS analyzed all-atom MD simulations. AS, JM and EF contributed to writing the article. EF supervised RS, AS and TGCB.

## Data and Software Availability

The three datasets, the MD simulations converted to the CG representation, the results obtained by iNMA and in-house scripts can be found in the repository Zenodo (doi: 10.5281/zenodo.7384244).

## Acknowledgement

The authors thank all members of the team “Structure et traduction des ARN viraux” at CiTCoM for useful comments and suggestions all along the process.

## Supporting Information Available

The Supporting Information is available free of charge on the ACS Publications website at DOI:

RMSF profiles; Figures on the analysis conducted on MD ensembles; Figures on the analysis performed on experimental ensembles; list of RNAs molecules studied (three dataset); Summary of the results obtained for the study of the unbound-to-transition.

