## Supplementary Information for "Internal Normal Mode Analysis applied to RNA flexibility and conformational changes"

### Supplementary Information for Internal Normal Mode Analysis for RNA flexibility and conformational changes

#### S1 RMSF profiles

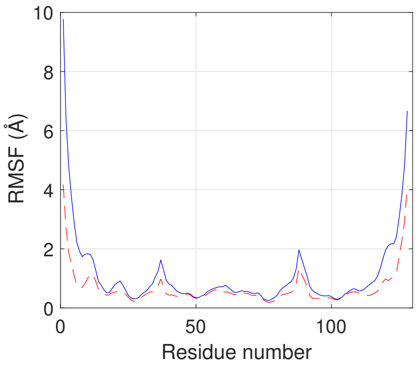


Figure S1: RMSF obtained for the PDB structure 1L9A using the first (red dashed line) and second (red) order approximation in eq. (12) and $R_c = 12$ Å and $\gamma = 0.6\,\mathrm{kcal\,mol^{-1}}$.

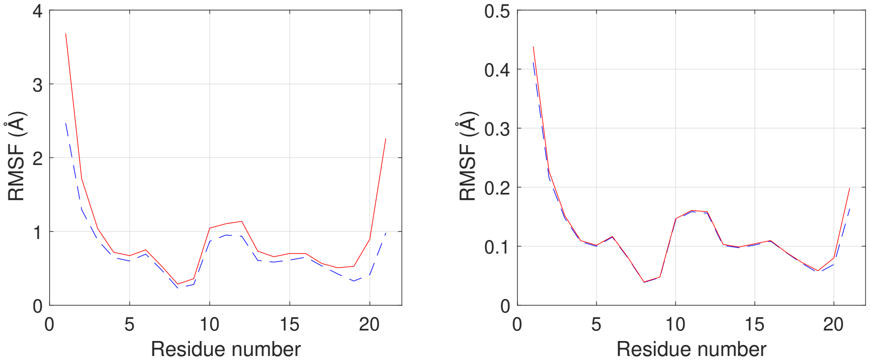


Figure S2: Left: RMSF obtained for the PDB structure 1QWA using the first (red dashed line) and second (red) order approximation in eq.( 12) and $R_c = 16$ Å and $\gamma = 0.1\,\mathrm{kcal\,mol^{-1}}$. Right: RMSF obtained for the PDB structure 1QWA using the first (red dashed line) and second (red) order approximation in eq. (12) and $R_c = 16$ Å and $\gamma = 0.6\,\mathrm{kcal\,mol^{-1}}$.

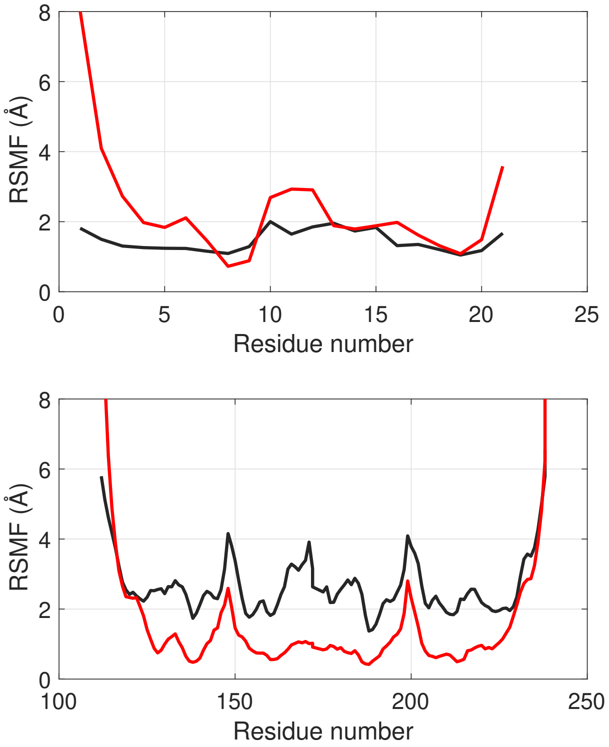


Figure S3: Comparison of iNMA RMSF (red) with RMSF from all-atom MD simulation (dark grey) computed for the PDB structure 1QWA (top) and 1L9A (bottom) using a value of distance cut-off $R_c = 16$ Å and a force constant $\gamma = 0.6\,\mathrm{kcal\,mol^{-1}}$. The RMSF profiles obtained via iNMA are multiplied by 18.

### S2 Figures for the analysis of the MD ensembles

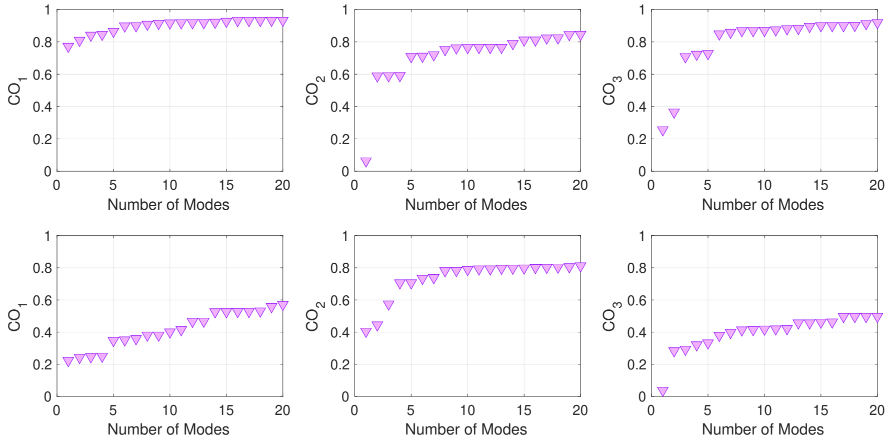


Figure S4: Cumulative overlap (CO) (see eq. (16)) as a function of the number of lowest modes used in the calculation for the first (left), the second (middle) and the third (right) principal components. In top, the results obtained for the PDB structure 2TRA are reported. In the bottom, the results for the PDB structure 1SDR. Red circle: HiRE-RNA model. Lilac downward-pointing triangle: RNA three-bead model. $R_c = 11\,\text{Å}$ for the RNA three-bead.

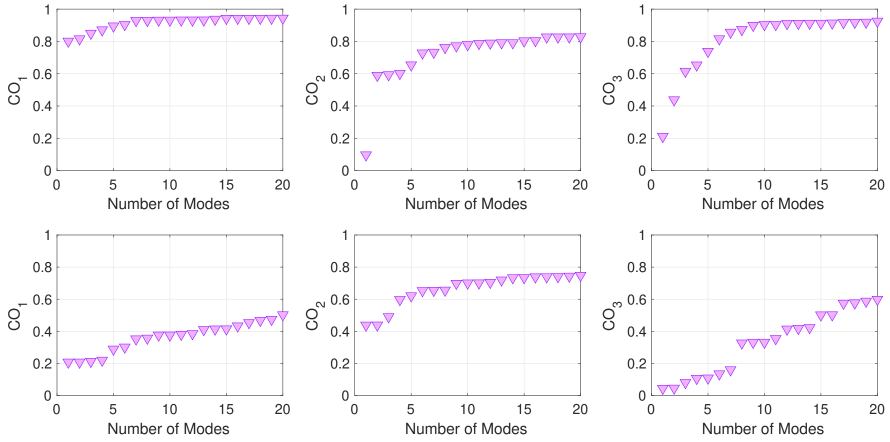


Figure S5: Cumulative overlap (CO) (see eq. (16)) as a function of the number of lowest modes used in the calculation for the first (left), the second (middle) and the third (right) principal components. On top, the results obtained for the PDB structure 2TRA are reported. In the bottom, the results for the PDB structure 1SDR. Red circle: HiRE-RNA model. Lilac downward-pointing triangle: RNA three-bead model. $R_c = 14\,\text{Å}$ for the RNA three-bead.

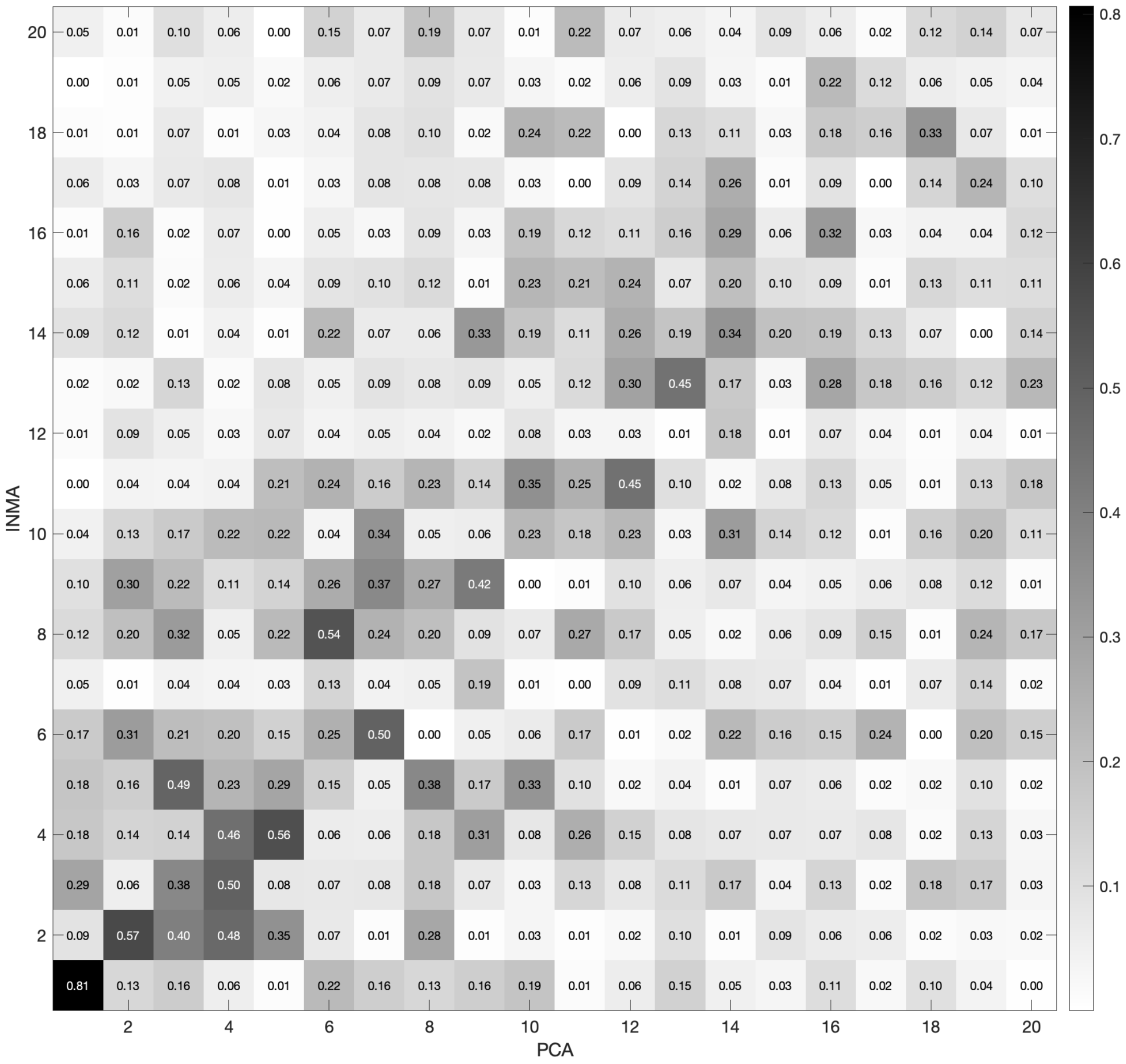


Figure S6: Matrix of overlap $O_{jk}$ between the internal mode $j$ and the principal component $k$ computed on the MD simulation of the PDB structure 2TRA. The distance cut-off $R_c$ is equal to 16 Å. The color of each cell is based on the calculated $O_{jk}$ and goes from white ($O_{jk}$ equal to zero) to black ($O_{jk}$ equal to 1).

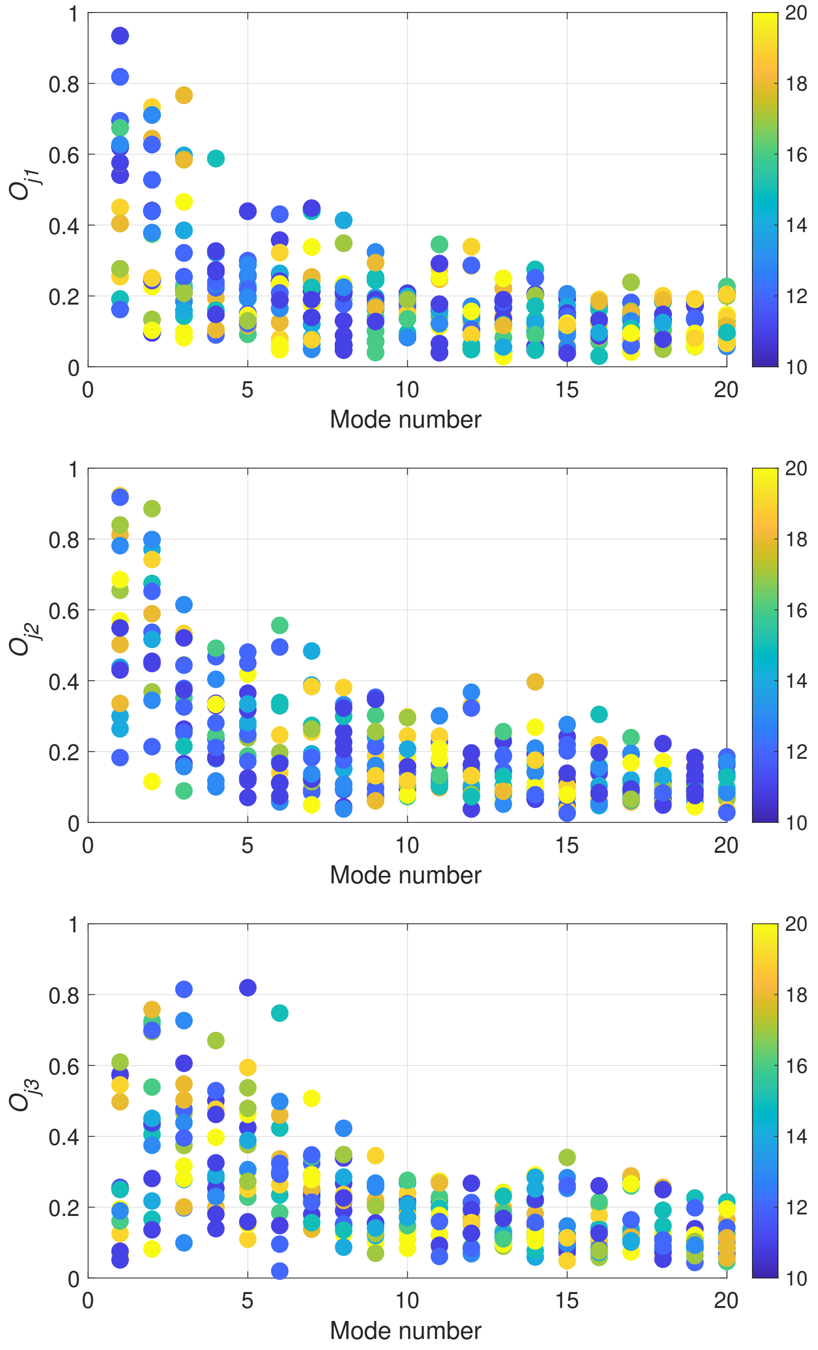


Figure S7: Maximum overlap $O_{j1}$ (top), $O_{j2}$ (middle) and $O_{j3}$ (bottom) for the lowest 20 internal modes. The dots are colored based on the value of the distance cut-off $R_c$ from blue to yellow. For example, $R_c = 16\,\text{Å}$ for green dots.

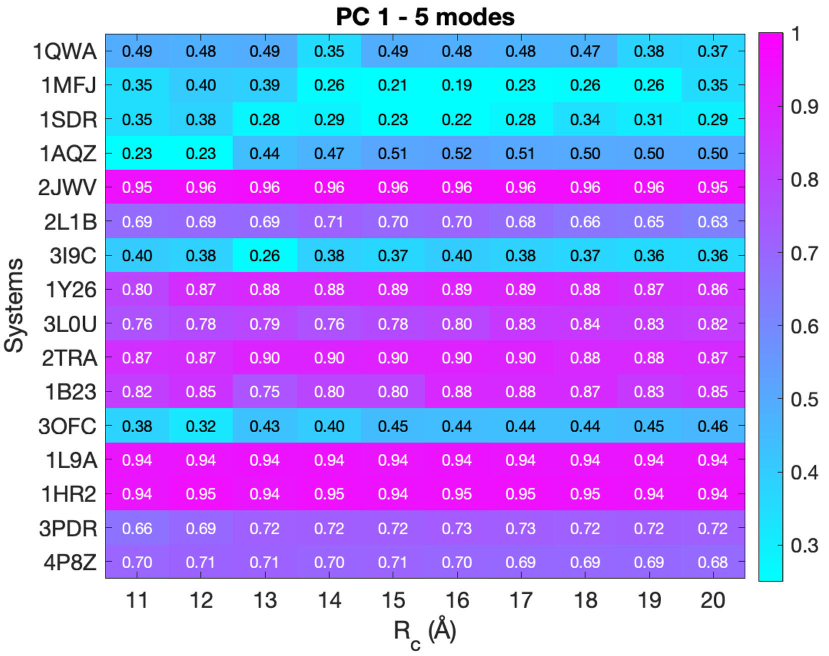


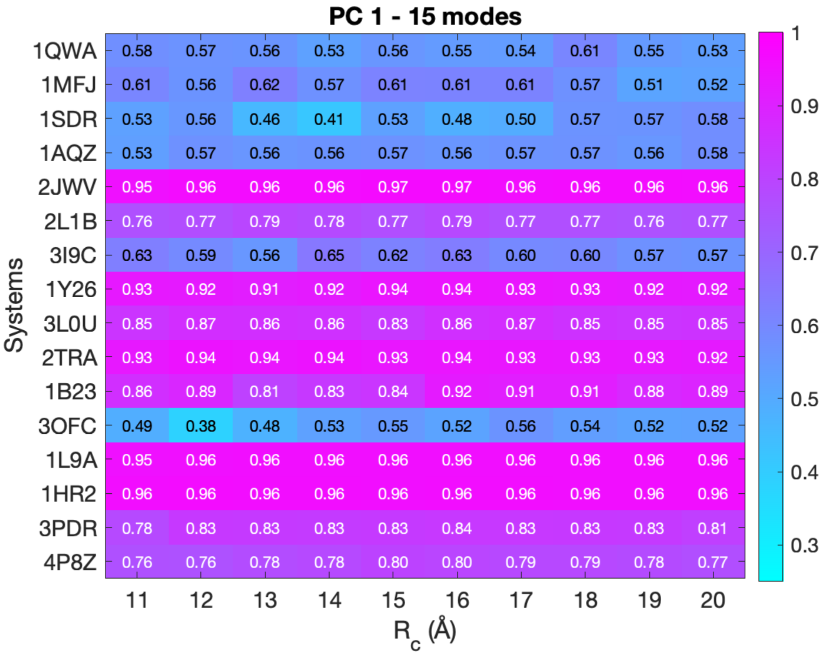


Figure S8: Cumulative overlap (CO) (see eq. (16)) computed using the five (top) and fifteen (bottom) low-frequency modes and the first principal component (PC1) for all the systems under investigation in dataset 1. The color of each cell is based on the calculated CO and goes from cyan (low CO, meaning that less than 30% of the motions are captured) to magenta (CO equal to 1, i.e., all the motions are captured and recovered).

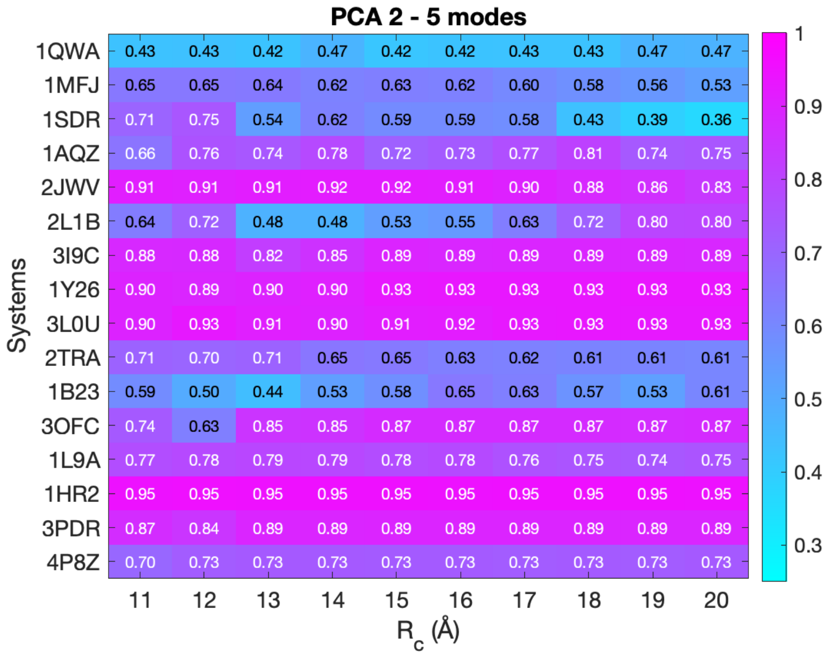


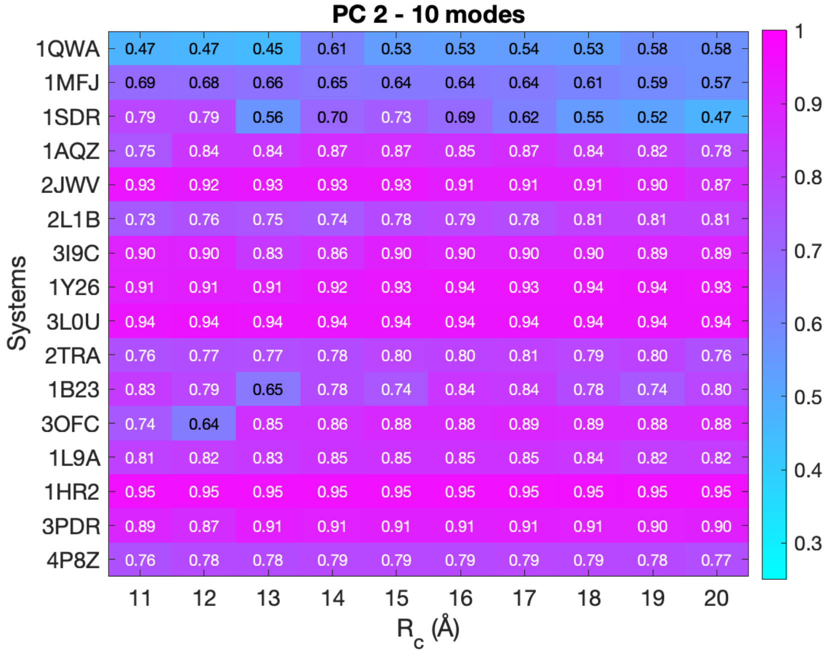


Figure S9: Cumulative overlap (CO) (see eq. (16)) computed using the first five (top) and ten (bottom) low-frequency modes and the second principal component (PC2) for all the systems under investigation in dataset 1. The color of each cell is based on the calculated CO and goes from cyan (low CO, meaning that less than 30% of the motions are captured) to magenta (CO equal to 1, i.e., all the motions are captured and recovered).

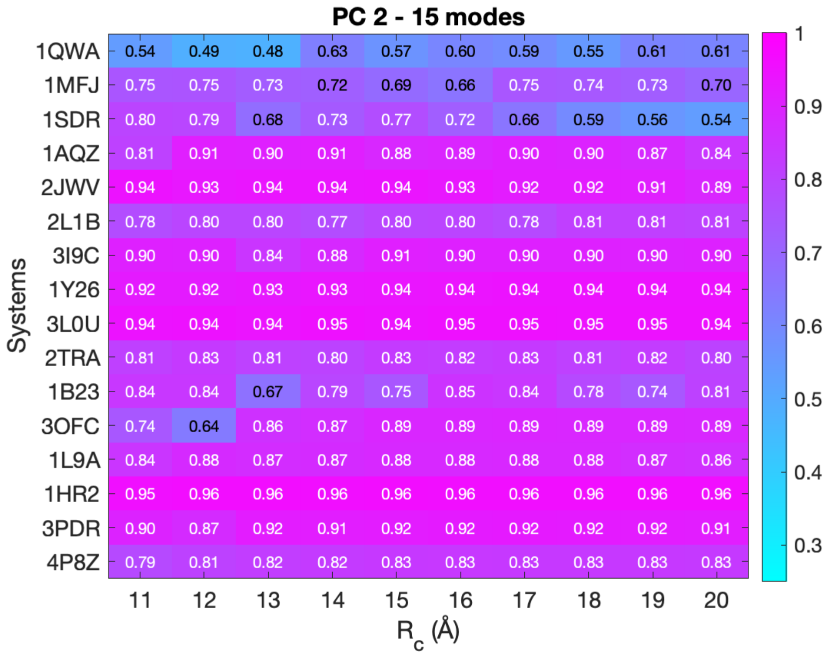


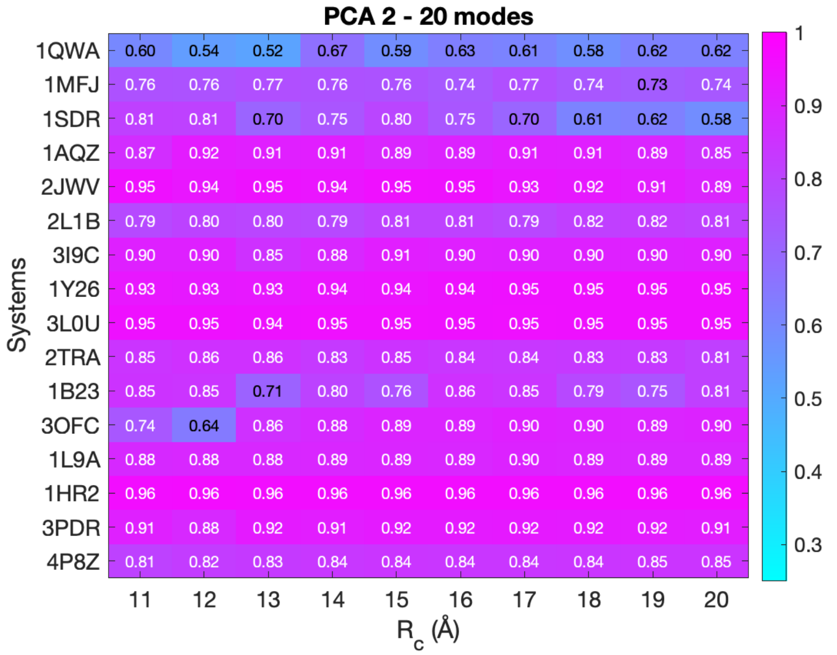


Figure S10: Cumulative overlap (CO) (see eq. (16)) computed using the fifteen (top) and twenty (bottom) low-frequency modes and the second principal component (PC2) for all the systems under investigation in dataset 1. The color of each cell is based on the calculated CO and goes from cyan (low CO, meaning that less than 30% of the motions are captured) to magenta (CO equal to 1, i.e., all the motions are captured and recovered).

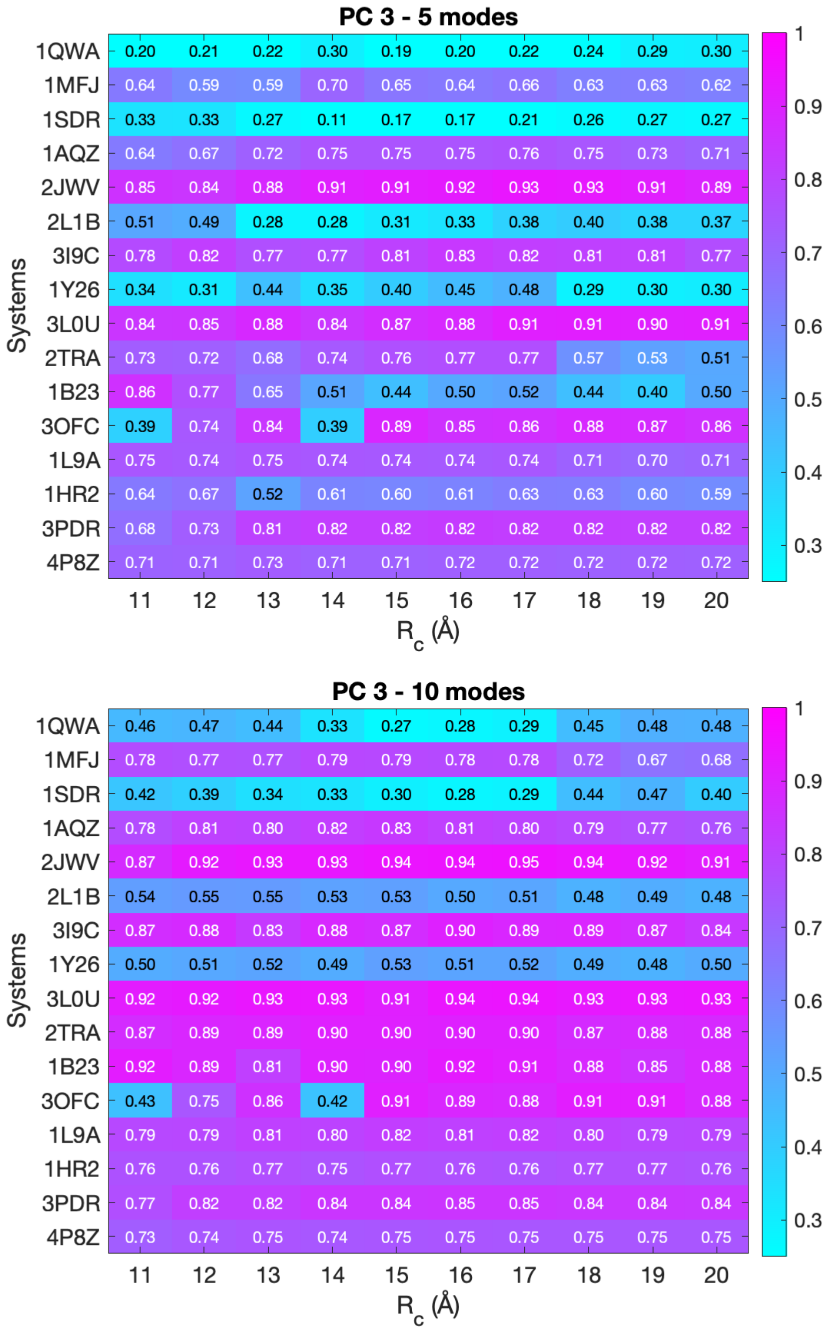


Figure S11: Cumulative overlap (CO) (see eq.( 16)) computed using the first five (top) and ten (bottom) low-frequency modes and the third principal component (PC3) for all the systems under investigation in dataset 1. The color of each cell is based on the calculated CO and goes from cyan (low CO, meaning that less than 30% of the motions are captured) to magenta (CO equal to 1, i.e., all the motions are captured and recovered).

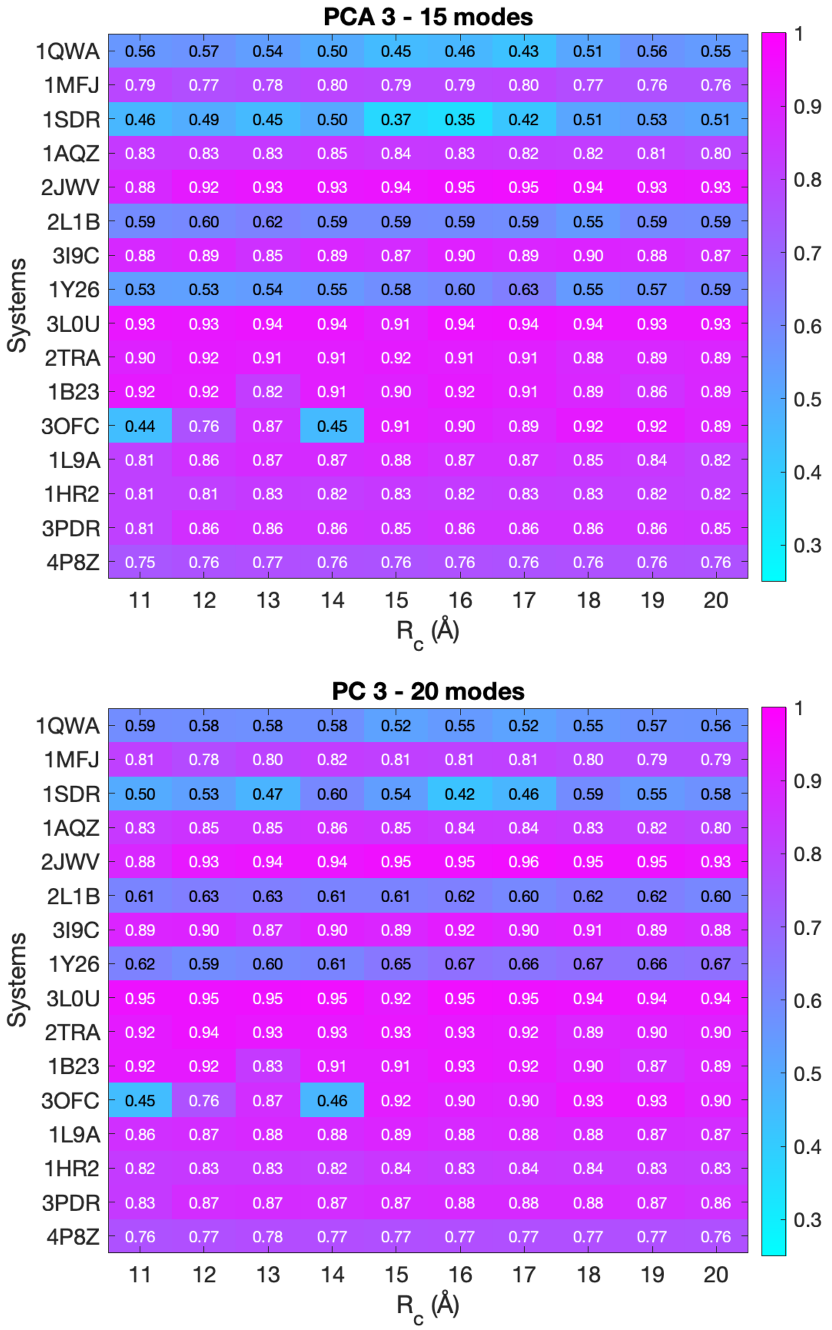


Figure S12: Cumulative overlap (CO) (see eq. (16)) computed using the fifteen (top) and twenty (bottom) low-frequency modes and the third principal component (P3) for all the systems under investigation in dataset 1. The color of each cell is based on the calculated CO and goes from cyan (low CO, meaning that less than 30% of the motions are captured) to magenta (CO equal to 1, i.e., all the motions are captured and recovered).

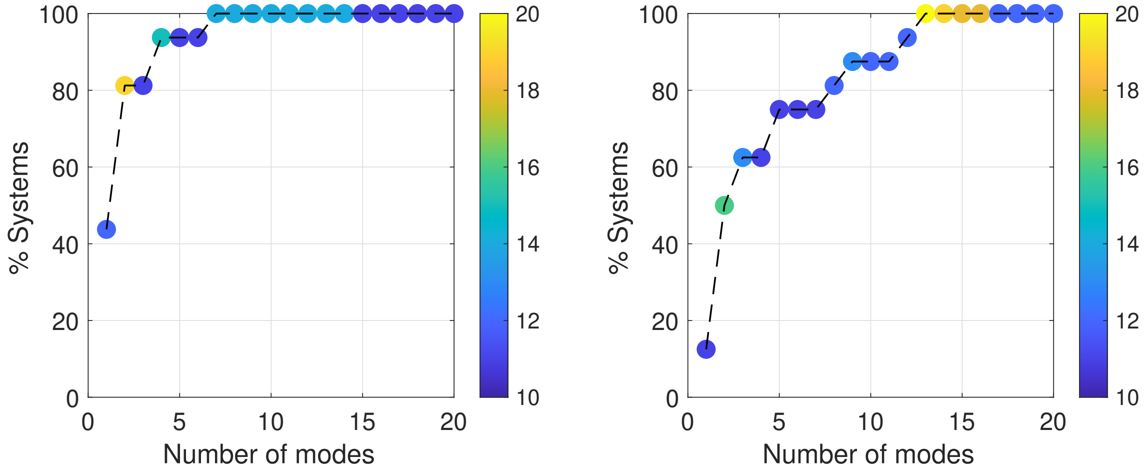


Figure S13: Percentage of systems of dataset 2 which capture at least 50% of the motions of the first principal component (left) and second one (right) as a function of the number of modes taken into account. The dots are colored based on the value of the distance cut-off $R_c$ from blue to yellow. For example, $R_c = 16\,\text{Å}$ for green dots.

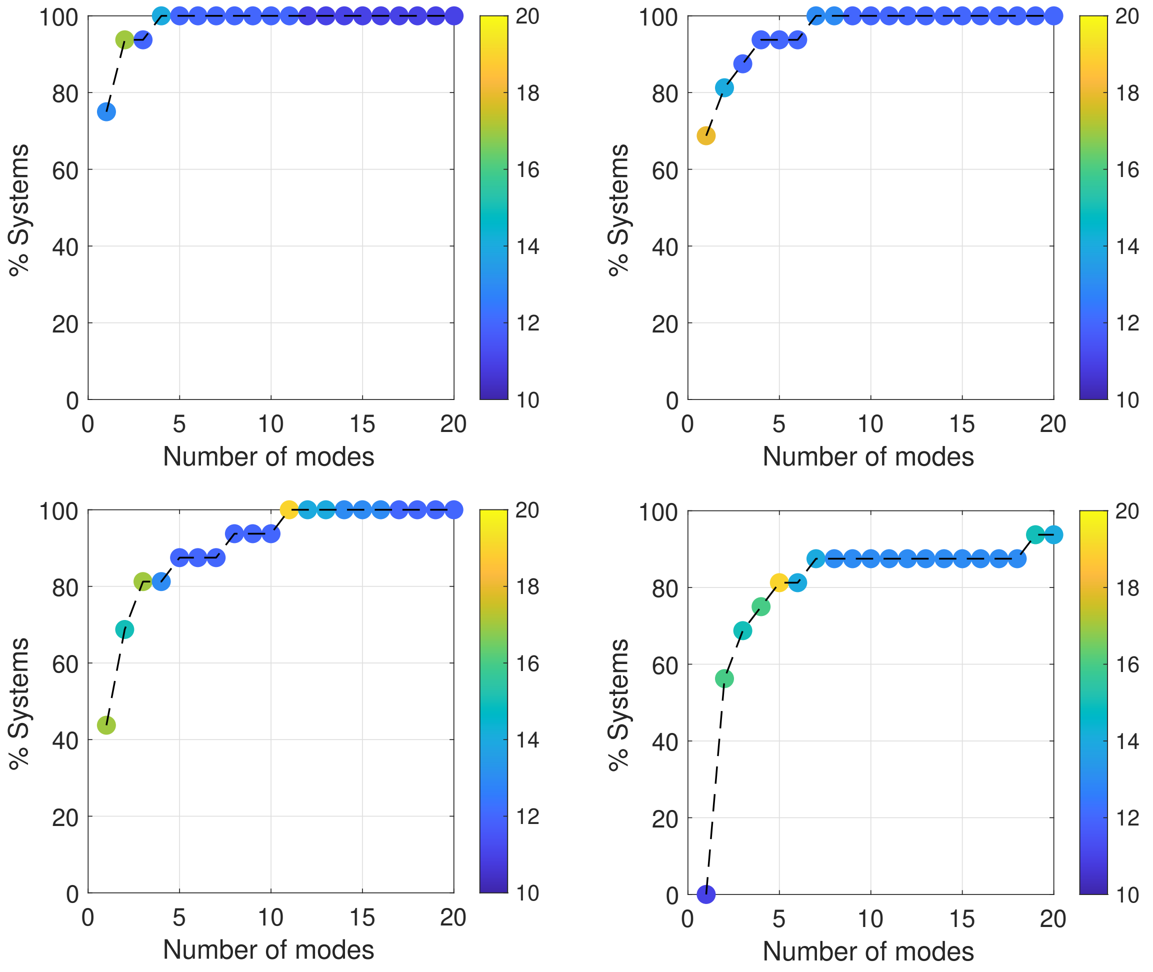


Figure S14: Percentage of systems of dataset 1 which capture at least 50% (top, left), 60% (top, right), 70% (bottom, left) and 80% (bottom, right) of the motions of three dominant PCs as a function of the number of modes taken into account (see eq. (18) for the definition of RMSIP). The dots are colored based on the value of the distance cut-off $R_c$ from blue to yellow. For example, $R_c = 16$ Å for green dots.

### S3 Figures for the analysis of experimental ensembles

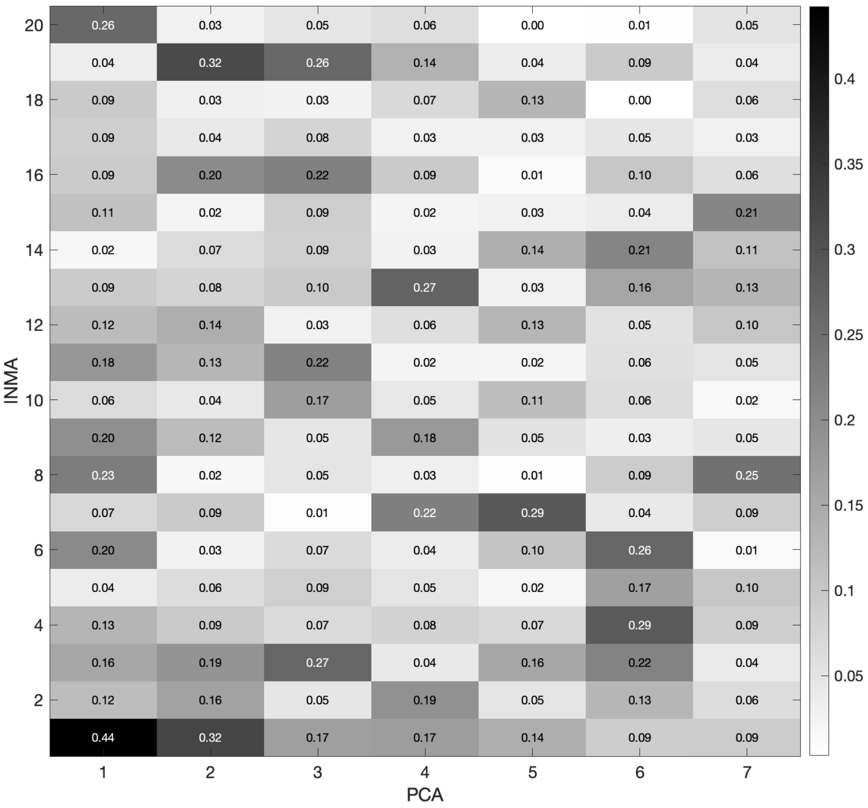


Figure S15: Matrix of overlap $O_{jk}$ between the internal mode $j$ and the principal component $k$ computed on the experimental ensemble of the U2 spliceosomal RNA (RF00004). The distance cut-off $R_c$ is equal to 16 Å. The color of each cell is based on the calculated $O_{jk}$ and goes from white ($O_{jk}$ equal to zero) to black ($O_{jk}$ equal to 1).

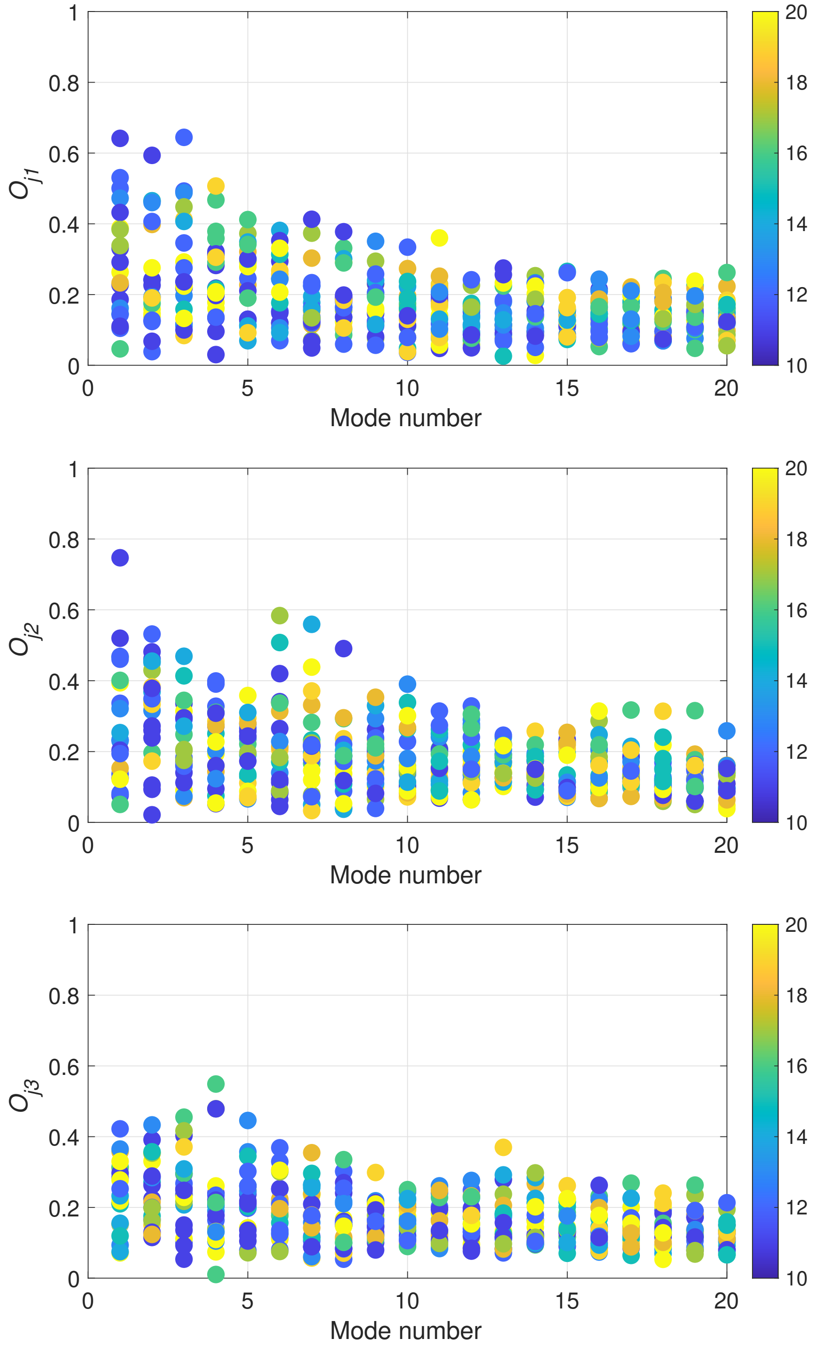


Figure S16: Maximum overlap $O_{j1}$ (top), $O_{j2}$ (middle) and $O_{j3}$ (bottom) for the lowest 20 internal modes of dataset 2. The dots are colored based on the value of the distance cut-off $R_c$ from blue to yellow. For example, $R_c = 16$ Å for green dots.

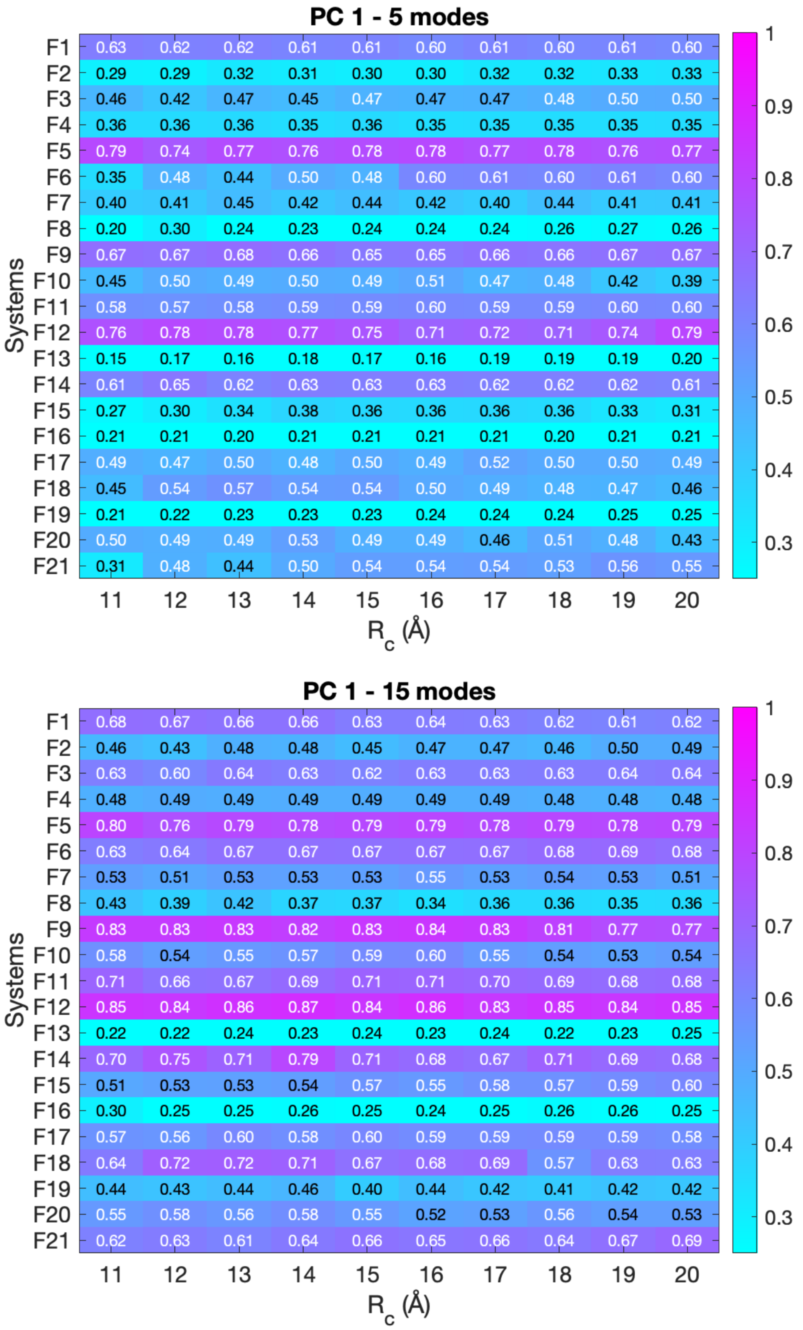


Figure S17: Cumulative overlap (CO) (see eq. (16)) computed using the five (top) and fifteen (bottom) low-frequency modes and the first principal component (PC1) for all the systems under investigation in dataset 2. The color of each cell is based on the calculated CO and goes from cyan (low CO, meaning that less than 30% of the motions are captured) to magenta (CO equal to 1, i.e., all the motions are captured and recovered).

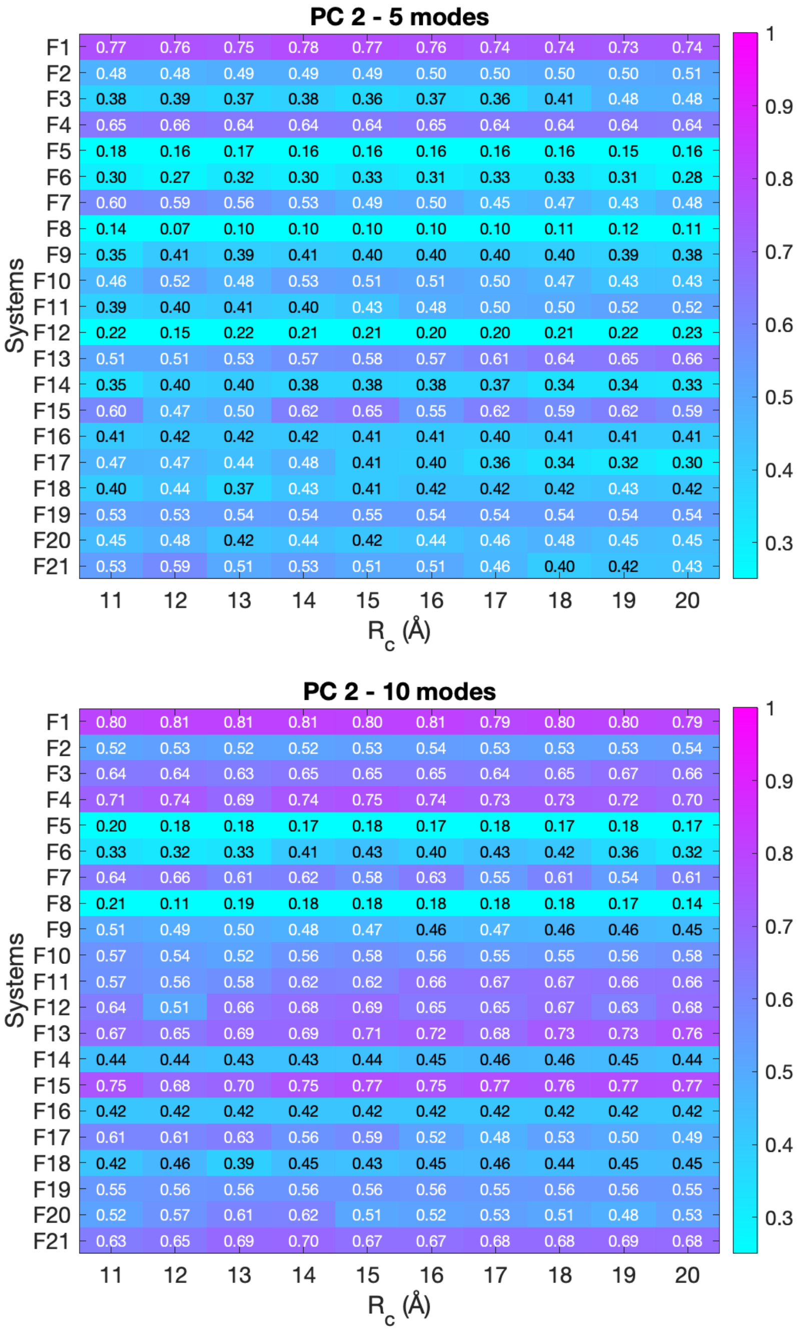


Figure S18: Cumulative overlap (CO) (see eq. (16)) computed using the first five (top) and ten (bottom) low-frequency modes and the second principal component (PC2) for all the systems under investigation in dataset 2. The color of each cell is based on the calculated CO and goes from cyan (low CO, meaning that less than 30% of the motions are captured) to magenta (CO equal to 1, i.e., all the motions are captured and recovered).

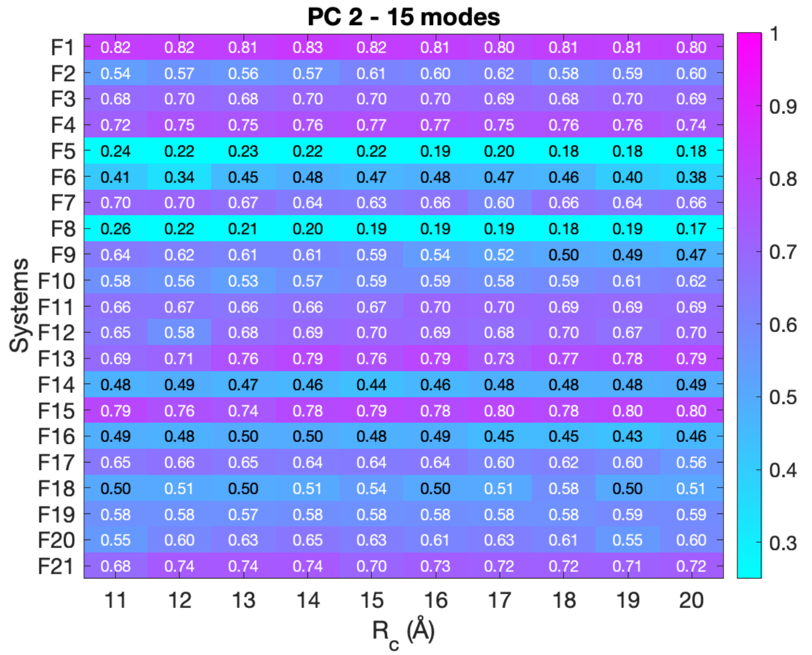


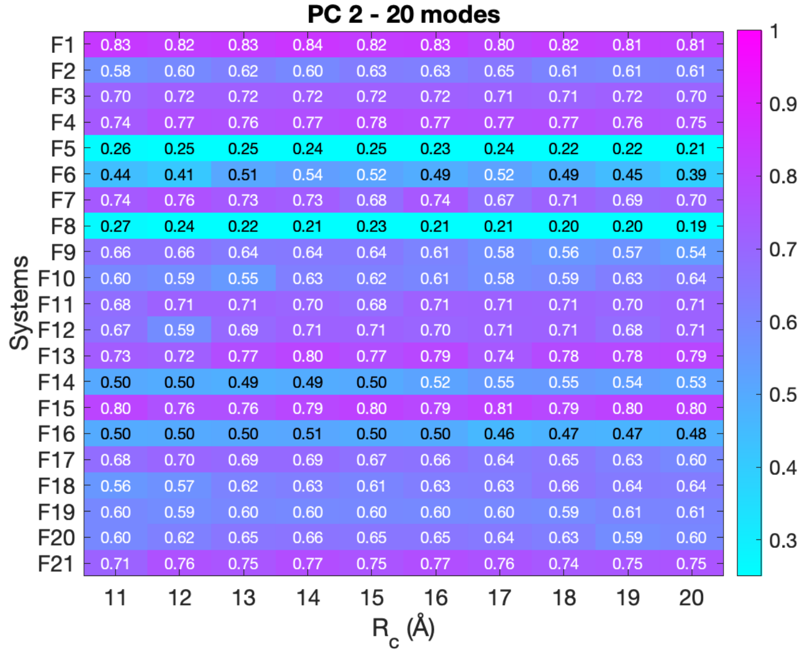


Figure S19: Cumulative overlap (CO) (see eq. (16)) computed using the fifteen (top) and twenty (bottom) low-frequency modes and the second principal component (PC2) for all the systems under investigation in dataset 2. The color of each cell is based on the calculated CO and goes from cyan (low CO, meaning that less than 30% of the motions are captured) to magenta (CO equal to 1, i.e., all the motions are captured and recovered).

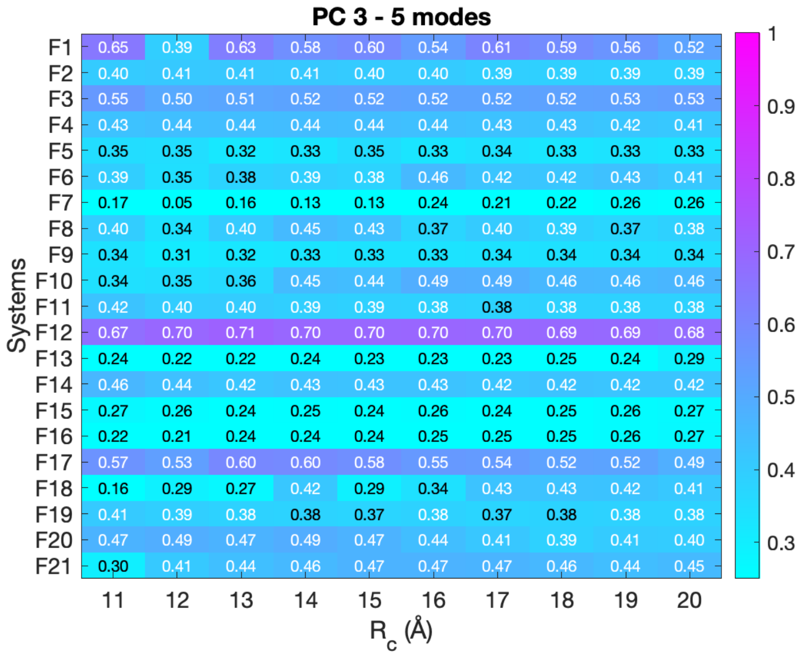


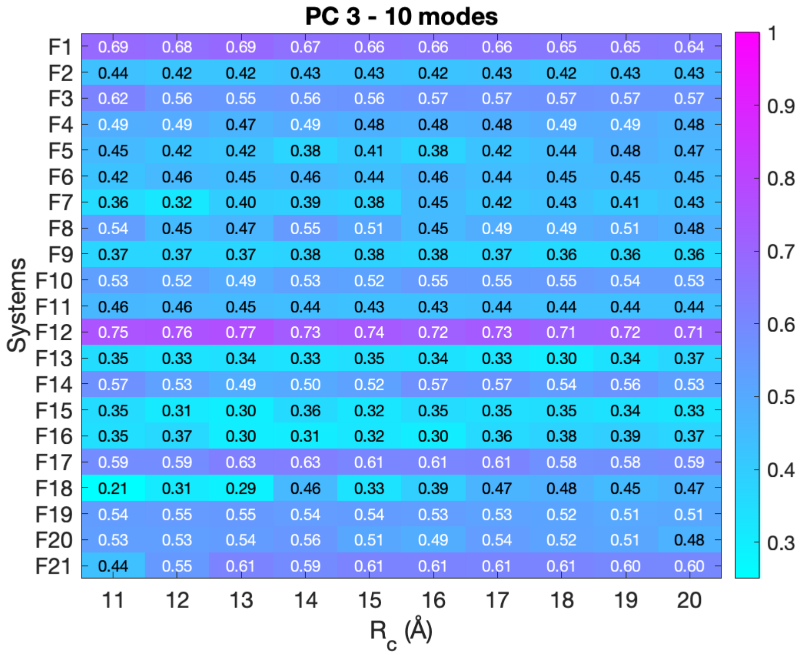


Figure S20: Cumulative overlap (CO) (see eq. (16)) computed using the first five (top) and ten (bottom) low-frequency modes and the third principal component (PC3) for all the systems under investigation in dataset 2. The color of each cell is based on the calculated CO and goes from cyan (low CO, meaning that less than 30% of the motions are captured) to magenta (CO equal to 1, i.e., all the motions are captured and recovered).

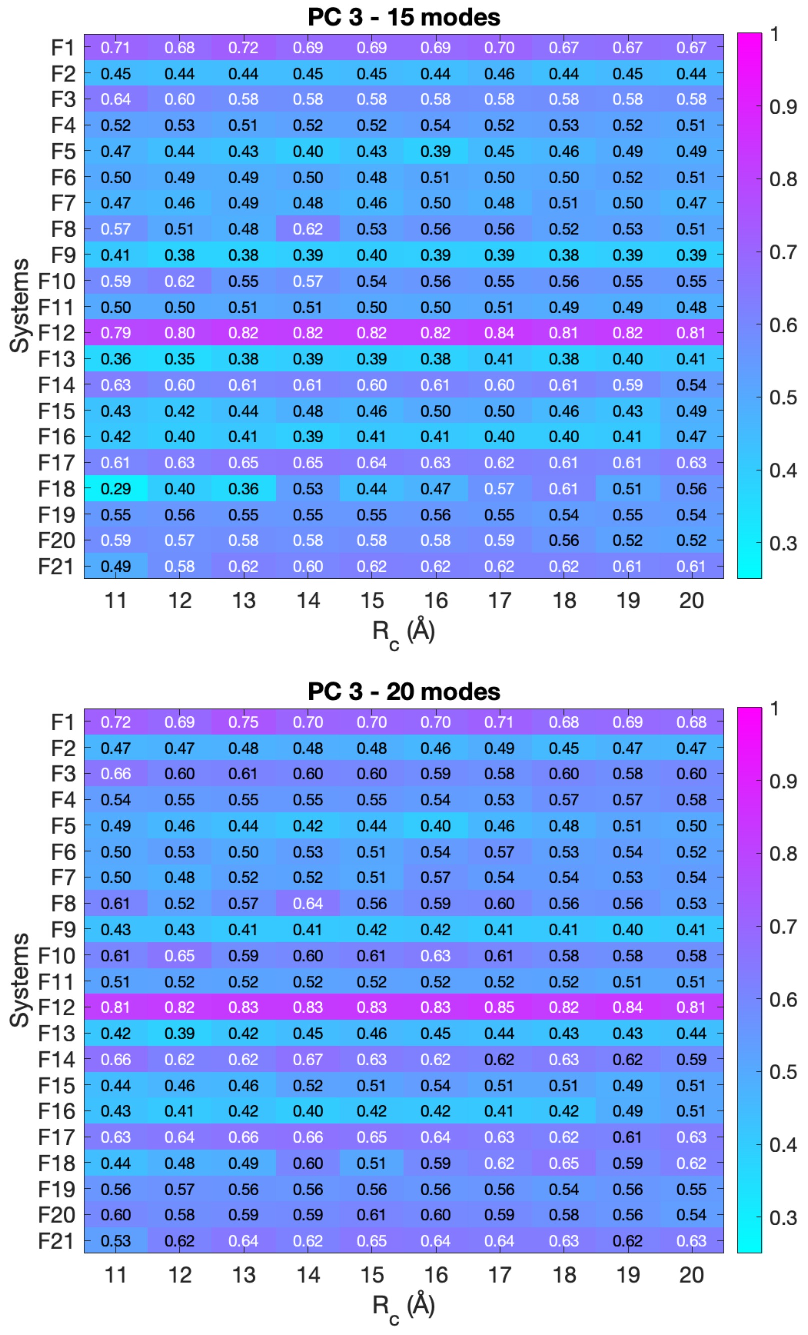


Figure S21: Cumulative overlap (CO) (see eq. (16)) computed using the fifteen (top) and twenty (bottom) low-frequency modes and the third principal component (PC3) for all the systems under investigation in dataset 2. The color of each cell is based on the calculated CO and goes from cyan (low CO, meaning that less than 30% of the motions are captured) to magenta (CO equal to 1, i.e., all the motions are captured and recovered).

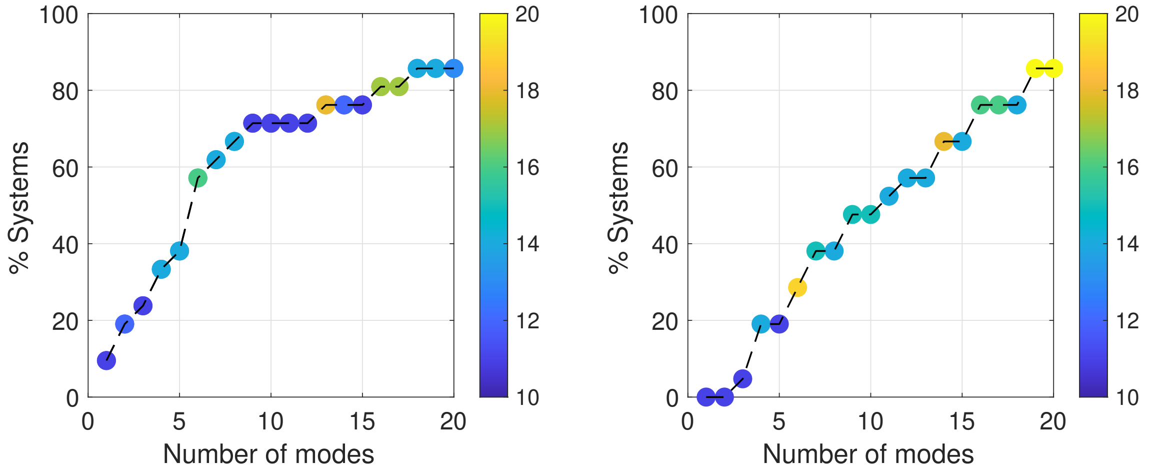


Figure S22: Percentage of systems of dataset 2 which capture at least 50% of the motions of the first principal component (left) and second one (right) as a function of the number of modes taken into account. The dots are colored based on the value of the distance cut-off $R_c$ from blue to yellow. For example, $R_c = 16$ Å for green dots.

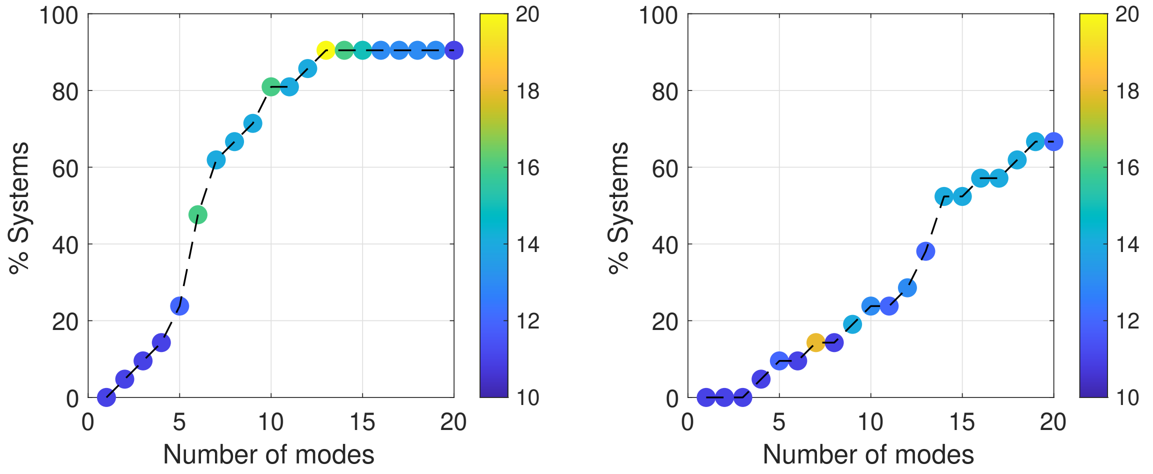


Figure S23: Percentage of systems of the dataset 2 which capture at least 50% (left) and 60% (right) of the motions of three dominant PCs as a function of the number of modes taken into account. The dots are colored based on the value of the distance cut-off $R_c$ from blue to yellow (see eq. (18) for the definition of RMSIP). For example, $R_c = 16$ Å for green dots.

### S4 Datasets

#### S4.1 Dataset 1 and 2

Table S1: Datasets 1 and 2. $N_b$: number of bases, $N_s$: number of structures. *: the cut-off was set at 0.3 Åto have more than two representative structures.

| ID | PDB ID | $N_b$ | ID | RFAM name | RFAM accession | $N_s$ | $N_b$ |
|---|---|---|---|---|---|---|---|
| A1 | 1QWA | 21 | F1 | tRNA | RF00005 | 40 | 73 |
| A2 | 1MFJ | 22 | F2 | 5.8sRNA | RF00002 | 37 | 150 |
| A3 | 1SDR | 23 | F3 | Metazoan SRP | RF00017 | 6 | 121 |
| A4 | 1AQZ | 27 | F4 | Group I catalytic intron | RF00028 | 5 | 238 |
| A5 | 2JWV | 28 | F5 | Group I catalytic intron | RF00028 | 6 | 157 |
| A6 | 2L1B | 34 | F6 | FMN riboswitch | RF00050 | 4 | 109 |
| A7 | 3I9C | 56 | F7 | TPP riboswitch | RF00059 | 4 | 53 |
| A8 | 1Y26 | 70 | F8 | HDV ribozyme | RF00094 | 7 | 44 |
| A9 | 3L0U | 72 | F9 | SAM riboswitch | RF00162 | 15 | 91 |
| A10 | 2TRA | 73 | F10 | Purine riboswitch | RF00167 | 6 | 60 |
| A11 | 1B23 | 76 | F11* | Lysine riboswitch | RF00168 | 5 | 173 |
| A12 | 3OFC | 119 | F12 | glmS riboswitch | RF00234 | 20 | 139 |
| A13 | 1L9A | 126 | F13* | glmS riboswitch | RF00234 | 6 | 120 |
| A14 | 1HR2 | 158 | F14 | Glycine riboswitch | RF00504 | 7 | 85 |
| A15 | 3PDR | 160 | F15 | GEMM cis-reg | RF01051 | 7 | 86 |
| A16 | 4P8Z | 188 | F16 | 23S rRNA pseudoknot | RF01118 | 34 | 108 |
| | | | F17 | Guanidine-III riboswitch | RF01763 | 4 | 41 |
| | | | F18 | U2 spliceosomal RNA | RF00004 | 8 | 43 |
| | | | F19 | 5SRNA | RF00001 | 74 | 118 |
| | | | F20 | U6 spliceosomal RNA | RF00026 | 9 | 44 |
| | | | F21 | U6 spliceosomal RNA | RF00026 | 11 | 102 |

##### S4.1.1 Database 2(RFarm)

- F1 (RF00005): 1EHZ_A (reference structure), 1TTT_D, 1TTT_E, 1TTT_F, 3DEG_A, 3IZY_N, 3WC2_P, 3WC2_, 4V4I_0, 4V4J_2, 4V4P_BB, 4V4P_BC, 4V4S_AW, 4V4W_AV, 4V7H_A7, 4V65_AA, 4V66_AA, 4V69_AY, 4YYE_C, 4YYE_D, 5M1J_A3, 6AH3_T, 6GQ1_AX, 6GQB_AX, 6GQV_AY, 6GZ5_Bw, 1I9V_A, 1SZ1_E, 4V4R_AW, 4V65_AE, 4V65_AP, 5MRC_bb, 5MRE_bb, 6GZ3_Bw, 4V4R_AV, 1JGP_B, 1MJ1_D, 1JGO_C, 1FCW_A, 1ZO1_F

- F2: 4U4R_8 (reference structure), 3J6X_8S, 3J6Y_8S, 6CB1_2, 6GQB_4, 6GQV_4, 6HD7_4, 6I7O_BS, 6I7O_YS, 3J77_8S, 3J78_8S, 3JCT_2, 4V7F_2, 5APO_8, 5I4L_4, 5I4L_8, 5JUS_C, 5JUT_C, 5JUU_C, 5LYB_4,5LYB_8, 5MEI_4, 5NDV_4, 5NDW_4, 5OBM_4, 6ELZ_2, 6EM3_2, 5IT7_8, 4V88_A4, 5DAT_4, 5H4P_4, 5NDW_8, 5NDV_8, 5TBW_4, 6EM4_2, 5Z3G_B
- F3: 1L9A_B (reference structure), 1RY1_A, 2J37_A, 4P3E_A, 5M73_A, 1MFQ_A
- F4: 1GRZ_B (reference structure), 1X8W_A, 1X8W_B, 1X8W_C, 1X8W_D
- F5: 1GID_B (reference structure), 1GID_A, 6D8M_A, 6D8O_A, 6D8O_B, 6BJX_A
- F6: 1L9A_B (reference structure), 1RY1_A, 2J37_A, 4P3E_A, 5M73_A, 1MFQ_A
- F7: 2CKY_A (reference structure), 3K0J_F, 3K0J_E, 2D2G_A
- F8: 1CX0_B (reference structure), 1VC6_B, 1VC7_B, 2OIH_B, 2OJ3_B, 1SJF_B, 1SJ3_R
- F9: 4b5r_A (reference structure), 5fkh_A, 5fk3_A, 5fk6_A, 5fk4_A, 2ydh_A, 3iqr_A, 3gx5_A, 4aob_A, 5fk5_A, 5fk2_A, 5fkg_A, 3gx7_A, 3gx3_A, 3iqn_A
- F10: 4FE5_B (reference structure), 1Y27_X, 3DS7_A, 3DS7_B, 3G4M_A, 3GAO_A
- F11: 3DIG_X (reference structure), 3DIO_X, 3DIS_A, 3DIY_A, 3DIL_A
- F12: 3G8T_Q (reference structure), 3l3c_R, 3l3c_Q, 3l3c_P, 3g9c_S, 3g9c_R, 3g9c_Q, 3g9c_P, 3g96_S, 3g96_P, 3g8s_P, 3g8s_S, 2nz4_P, 2nz4_Q, 2nz4_R, 2nz4_S, 3l3c_S, 3g8s_R, 3g8t_P, 3g96_R
- F13: 2Z75_B (reference structure), 3B4B_B, 4MEG_B, 2H0S_B, 2HO7_B, 2GCV_B
- F14: 3OXJ_A (reference structure), 3OWZ_A, 3OWZ_B, 3OXD_B, 3OXJ_B, 3OX0_B, 3OXE_B

- F15: 3MXH_R (reference structure), 3UCZ_R, 3MUV_R, 3MUT_R, 3UD4_R, 3UCU_R, 3MUR_R
- F16: We refined the ensemble RF01118_g1_v3.MSA.pdb provided as supplementary materials in.[1]
- F17: 5NWQ_A (reference structure), 5NZ6_A, 5NZD_A, 5NWQ_B
- F18: 5gmk_L (reference structure), 5gm6_L, 5ylz_F , 5lqw_2, 5wsg_L, 5y88_F , 5lj3_Z, 5mps_2,
- F19: 5J8B_B (reference structure), 4V8E_AB, 4V8F_DB, 4V8J_BB, 4V8N_DB, 4V8Q_AB, 1Y69_9, 4W4G_RB, 4W4G_YB, 3DLL_Z, 4WT8_Cs, 4WT8_Ds, 1SM1_9, 3FWO_B, 4XEJ_A5, 4XEJ_B5, 3JQ4_B, 5EL5_1J, 5IMQ_E, 5IMR_E, 5OT7_5, 5ZLU_X, 4LEL_YB, 6C5L_BB, 6GZQ_B1, 6GZX_B1, 6GZX_B2, 6GZZ_B1, 6GZZ_B2, 6N9E_2B, 4V4I_x, 4V4J_x, 4V5F_BB, 4V5K_BB, 4V5S_BB, 4V63_BB, 4V63_DB, 4V67_DB, 4V6G_BB, 1NKW_9, 4V8U_BB, 4V8X_BB, 4V9Q_CB, 3PIO_Y, 5A9Z_AB, 5DM7_Y, 5UQ7_B, 6BOH_C, 4V7P_BB, 4V8H_DB, 4V9J_DB, 4Y4P_2B, 4IOC_Y, 4V5E_DB, 4V5J_BB, 6C5L_DB, 4V8G_BB, 4V5R_BB, 4V90_BB, 4V9J_BB, 4V51_BB, 4V9L_DB, 2ZJP_Z, 4WFN_Y, 5D8B_AD, 4WSM_16, 5EL5_16, 4WQR_16, 6B4V_GB, 6BOK_C, 4V9S_BB, 4V9K_DB, 5VPP_RB, 4V5C_DB
- F20: 6ID1_F (reference structure), 5MQF_6, 6QDV_6, 5XJC_F, 5YZG_F, 5Z58_F, 6ICZ_F, 6ID0_F, 6FF4_6
- F21: 5LJ3_V (reference structure), 5GM6_E, 5YLZ_D, 6BK8_6, 6EXN_6, 5LQW_6, 5WSG_E, 6J6G_E, 6J6N_E, 5GMK_E, 5MPS_6

#### S4.2 Dataset of unbound-to-bound transitions

Table S2: Dataset of unbount-to-bound transition.u: unbound, b: bound, $RMSD_{exp}$: RMSD between the unbound and bound structure.*: different NMR structures.

| ID | PDB $ID_u$ | PDB $ID_b$ | $RMSD_{\mathrm{exp}}$ (Å) |
|---|---|---|---|
| UB1 | 1DUH | 1HQ1 | 37.1 |
| UB2 | 2ZUF | 1J2B | 11.0 |
| UB3 | 1JZX | 2BH2 | 9.3 |
| UB4 | 1SDR | 3IEV | 5.7 |
| UB5 | 1B23 | 1U0B | 6.9 |
| UB6 | 1U0B | 1B23 | 6.9 |
| UB7 | 2JWV* | 1OOA | 5.3 |
| UB8 | 2JWV* | 1OOA | 5.5 |
| UB9 | 2JWV* | 1OOA | 5.1 |
| UB10 | 3TRA | 1ASY | 5.0 |
| UB11 | 1QWA | 1RKJ | 5.2 |
| UB12 | 1EHZ | 1R3E | 2.8 |
| UB13 | 2TRA | 1ASY | 5.3 |
| UB14 | 1U63 | 2VPL | 4.1 |
| UB15 | 1JU7 | 1ZBH | 4.3 |
| UB16 | 3CW5_a | 2FMT | 3.4 |
| UB17 | 2AZX | 2AKE | 3.9 |
| UB18 | 3CW5_A | 2FMT | 3.4 |
| UB19 | 1EFW | 1C0A | 2.2 |
| UB20 | 1UN6_E | 1UN6_F | 2.5 |
| UB21 | 1MFK | 2PJP | 3.1 |
| UB22 | 2B7G | 2B6G | 2.3 |
| UB23 | 3CW6 | 2FMT | 3.4 |
| UB24 | 2V3C | 1LNG | 2.0 |

### S5 Results on the study of unbound-to-bound transitions

Table S3: Predicting the conformation change for RNAs using a single iNMA mode. The following information is given: PDB ID code for the unbound structures, best mode based on the overlap or RMSD, the final RMSD ($RMSD_{m-b}$) and MSD between the unbound and bound structures ($RMSD_{exp}$).

| ID | Best overlap mode | Best RMSD mode | $RMSD_{m-b}$ (Å) | $RMSD_{exp}$ (Å) |
|---|---|---|---|---|
| UB1 | 4 | 16 | 11.2 | 37.1 |
| UB2 | 2 | 19 | 10.8 | 11.0 |
| UB3 | 1 | 16 | 7.4 | 9.3 |
| UB4 | 11 | 14 | 4.0 | 5.7 |
| UB5 | 1 | 2 | 4.0 | 6.9 |
| UB6 | 2 | 2 | 4.0 | 6.9 |
| UB7 | 5 | 2 | 4.3 | 5.3 |
| UB8 | 1 | 2 | 4.2 | 5.5 |
| UB9 | 1 | 1 | 4.0 | 5.1 |
| UB10 | 3 | 1 | 3.1 | 5.0 |
| UB11 | 6 | 1 | 4.5 | 5.2 |
| UB12 | 8 | 4 | 2.7 | 2.8 |
| UB13 | 1 | 1 | 3.2 | 5.3 |
| UB14 | 3 | 1 | 3.6 | 4.1 |
| UB15 | 18 | 1 | 3.9 | 4.3 |
| UB16 | 1 | 2 | 2.2 | 3.4 |
| UB17 | 19 | 17 | 3.8 | 3.9 |
| UB18 | 1 | 2 | 2.2 | 3.4 |
| UB19 | 6 | 6 | 2.1 | 2.2 |
| UB20 | 5 | 1 | 1.8 | 2.5 |
| UB21 | 1 | 1 | 2.8 | 3.1 |
| UB22 | 19 | 3 | 2.2 | 2.3 |
| UB23 | 1 | 2 | 2.2 | 3.4 |
| UB24 | 12 | 3 | 1.9 | 2.0 |
